# Multifunctional biomimetic porphyrin-lipid nanoparticles – novel nanoscale theranostics for atherosclerotic cardiovascular disease

**DOI:** 10.1101/2024.02.13.580218

**Authors:** Victoria Nankivell, Lauren Sandeman, Liam Stretton, Achini K. Vidanapathirana, Maneesha A. Rajora, Juan Chen, William Tieu, Peter J. Psaltis, Joanne T. M. Tan, Yung-Chih Chen, Karlheinz Peter, Gang Zheng, Christina Bursill

**Author notes:** Correspondence to: A/ Prof Christina Bursill Adelaide Medical School, Faculty of Health and Medical Science, University of Adelaide and the Vascular Research Centre, Lifelong Health Theme, South Australian Health and Medical Research Institute.

## Abstract

**Background:** High-density lipoprotein (HDL) nanoagents have unrealized potential for atherosclerosis theranostics. Porphyrin-lipid HDL mimetic nanoparticles (Por-HDL-NPs) incorporate porphyrin-lipid which permits near infrared fluorescence imaging and positron emission tomography (PET) through chelation of Copper-64 (^64^Cu). The outer shell contains apolipoprotein A-I mimetic peptide R4F that interacts with scavenger receptor SR-BI, enabling macrophage targeting and therapeutic effects. We leveraged the theranostic properties of Por-HDL-NPs for testing in atherosclerosis.

**Methods and Results:** *In vitro*, Por-HDL-NPs were internalised by immortalised bone marrow-derived macrophages (iBMDMs), visualised via fluorescence microscopy and flow cytometry. Por-HDL-NPs increased cholesterol efflux from [^3^H]-cholesterol-loaded iBMDMs, (49%, *P*<0.05), compared to reconstituted HDL. Incubation of iBMDMs with Por-HDL-NPs reduced mRNA levels of inflammatory mediators *Il-1β* (88%), *Il-18* (54%) and *Ccl5* (75%), and protein secretion of IL-1β (69%) and CCL5 (82%), *P*<0.05. Por-HDL-NPs suppressed inflammasome components *Nlrp3* (69%) and *Asc* (36%), *P*<0.05. Studies using siRNA deletion of SR-B1 and methyl-β-cyclodextrin, revealed the anti-inflammatory properties of Por-HDL-NPs were independent of SR-B1 and cholesterol efflux. However, Por-HDL-NPs suppressed activation of inflammatory transcription factor NF-κB (53%, *P*<0.05). In *Apoe*^-/-^ mice, PET imaging showed ^64^Cu-Por-HDL-NPs localised in hearts and detected increases in plaque over time with high-cholesterol diet. Por-HDL-NP fluorescence was visualised in aortic sinus plaques, co-localised with CD68^+^ macrophages, and by fluorescence IVIS imaging in aortic arch plaque. Por-HDL-NP-treated mice had smaller early-stage (22%) and unstable plaques (52%) and fewer circulating monocytes (32%) than control PBS-treated mice, *P*<0.05 for all.

**Conclusions:** Por-HDL-NPs have theranostic properties, exhibiting both multi-modal imaging capabilities for identifying plaque and athero-protective therapeutic effects.

**Clinical Perspective:** *What is new?:*

- Porphyrin high-density lipoprotein (HDL) mimetic nanoparticles (Por-HDL-NPs) have theranostic application in atherosclerosis.
- Por-HDL-NPs are internalized by macrophages *in vitro* and plaque macrophages *in vivo*, enabling the visualization of atherosclerosis by both positron emission tomography and multiple fluorescence imaging modalities.
- Por-HDL-NPs exhibit atheroprotective effects and suppress inflammation, promote cholesterol efflux, reduce atherosclerotic plaque development and lower the number of circulating monocytes.

*What are the clinical implications?:*

- The PET imaging and plaque targeting capabilities of Por-HDL-NPs have implications for improved non-invasive tracking of human atherosclerosis development.
- The excellent fluorescence imaging and plaque targeting properties of Por-HDL-NPs have clinical significance for improved detection of early-stage plaque using intravascular imaging strategies.
- Por-HDL-NPs provide therapeutic capabilities that target plaque directly, independent of lipid-lowering, suggestive of their potential to provide benefit on top of current lipid-lowering strategies.

## Introduction

Atherosclerosis is driven by inflammation and lipid accumulation in the affected arteries^1^ Despite substantial improvements in evidence-based preventative approaches, there remains an unacceptably high number of people who suffer cardiovascular complications as a result of atherosclerotic occlusions^2^. This illustrates the need for novel therapies and diagnostics that improve the management of atherosclerotic cardiovascular disease (CVD) into the future.

Nanoparticles are an exciting technology that can be designed to target atherosclerotic plaques and provide diagnostic or therapeutic effects^3–5^. Their greatest potential lies in strategies that combine diagnostic and therapeutic utility into one particle. A nanoparticle with the ability to enable multimodal plaque imaging and elicit pleiotropic atheroprotective actions including cholesterol efflux and the inhibition of inflammation, holds promise as an effective theranostic for atherosclerosis^6^.

Reconstituted high-density lipoprotein (rHDL)-like nanoparticles have been tested as theranostic agents for atherosclerosis^3,7–9^. They track to atherosclerotic plaques and can be detected by multiple imaging modalities. HDL-like nanoparticles offer therapeutic effects, reducing plaque size and suppressing plaque macrophage proliferation in murine models of atherosclerosis^3^. Into a phospholipid-based outer surface, these nanoparticles incorporate full-length apolipoprotein (apo)A-I, the main protein component of HDL that promotes cholesterol efflux, suppresses inflammation^10,11^ and facilitates targeting to plaque macrophages. To enable diagnostic properties,

HDL-like nanoparticles can be integrated with gadolinium for magnetic resonance imaging (MRI)^3^, the radioisotope zirconium-89 (^89^Zr) for positron emission tomography (PET) imaging^9^ or DiO/DiR for fluorescence imaging^3^. HDL-like nanoparticles have been loaded with statin drugs to extend their therapeutic benefits^3^. Overall, HDL-like nanoparticles offer highly desirable traits for successful theranostic application in atherosclerosis.

Porphyrin HDL mimetic nanoparticles (Por-HDL-NPs) are an HDL-like nanoparticle with multimodal imaging and therapeutic capabilities. Por-HDL-NPs contain porphyrin-lipid conjugates in their lipid shell simultaneously enabling excellent near-infrared fluorescence and non-invasive PET imaging capabilities through chelation of radionuclides such as Copper-64 (^64^Cu)^12–14^. Por-HDL-NPs also incorporate an apoA-I mimetic peptide ‘R4F’ (Ac-FAEKFKEAVKDYFAKFWD). R4F constrains the lipid shell to ∼20 nm in diameter via an α-helix network structure that facilitates cellular uptake^15^. Importantly, R4F allows for cell targeting and therapeutic capabilities via pathways that have benefit for atherosclerosis applications. The R4F structure interacts with the scavenger receptor class B type I (SR-BI), which is highly expressed on atherosclerotic plaque macrophages and mediates cell signaling events associated with cholesterol efflux/influx and inflammation^16–18^. Whilst previously explored for targeted cancer therapy^15,19^, Por-HDL-NPs have intriguing potential for atherosclerosis.

Accordingly, this study tested the ability of Por-HDL-NPs to detect and reduce atherosclerosis. *In vitro*, we found Por-HDL-NPs are internalized by macrophages, promote cholesterol efflux and inhibit inflammation. *In vivo*, infusions of Por-HDL-NPs in atherosclerosis-prone apolipoprotein (*Apo)e*^-/-^ mice fed a high-cholesterol diet were found to track to atherosclerotic plaques, visualized using fluorescence and PET imaging. Mice injected with Por-HDL-NPs developed less plaque, demonstrating atheroprotective properties. These findings have implications for the simultaneous identification and prevention of atherosclerosis using multifunctional Por-HDL-NP nanoscale theranostics.

## Methods

### Porphyrin HDL mimetic nanoparticles

The discoidal and cholesterol oleate (CO)-loaded porphyrin HDL mimetic nanoparticles (Por-HDL-NPs) were synthesized and characterized as per Supplemental Methods for nanoparticle synthesis. Table S1 summarizes nanoparticle compositions.

### 64Cu-labelling of Por-HDL-NPs

^64^Cu-Cl_2_ production and nanoparticle labelling were conducted by the Molecular Imaging and Therapy Research Unit (MITRU, SAHMRI, Adelaide). Por-HDL-NPs were labelled as previously described^13,15^. Radiochemical purity was verified by thin layer chromatography and activity measured prior to intravenous injection.

### Por-HDL-NP fluorescence detection in iBMDMs

Immortalized murine bone marrow derived macrophages (iBMDMs) were incubated with PBS or discoidal Por-HDL-NPs (10 µg/mL, porphyrin-lipid concentration) for 24h. Por-HDL-NP uptake was analyzed by flow cytometry on a BD LSRFortessa™ X-20 Analyzer (Becton Dickinson) with the red laser (excitation λ: 640 nm; emission λ: 670/30 nm) to detect porphyrin-lipid fluorescence. iBMDMs on microscope slides were mounted with VECTASHIELD® antifade medium with DAPI (Vector Laboratories) and porphyrin-lipid fluorescence detected by confocal microscopy (Leica, excitation λ: 653 nm; emission λ: 670 nm beyond).

### Cholesterol efflux

iBMDMs were incubated with 0.074 MBq/mL of [1,2-3H(N)]-labelled cholesterol (Perkin-Elmer) for 24h at 37°C, then with PBS, reconstituted HDL (rHDL, 25 µg/mL, ApoA-I concentration), and discoidal or CO-loaded Por-HDL-NPs (10 or 25 µg/mL, R4F concentration) for 4h. Cholesterol efflux was determined as % counts per minute (CPM) in the supernatant versus cell lysates with a liquid scintillation analyzer (Tricarb-2810TR, Perkin Elmer).

### *In vitro* inflammation

iBMDMs were incubated with PBS, discoidal or CO-loaded Por-HDL-NPs (10 µg/mL) for 18h and then stimulated with lipopolysaccharide (LPS, 10 ng/mL, L4391 Sigma) or murine interferon (IFN)-γ (10 ng/mL, 315-05 Peprotech) for 16h.

### Real time-qPCR

RNA was extracted from treated iBMDMs and murine aortic arches with TRI-reagent (Sigma-Aldrich) then reverse transcribed to cDNA using iScript Reverse Transcriptase Supermix (Biorad). Primers for inflammatory and housekeeper genes (Table S2) measured gene expression changes by qPCR, calculated using the ^ΔΔ^*Ct* method referenced to housekeeper genes *Rplp0* or *Beta-2 microglobulin* (*B2m)*.

### ELISAs

Secreted proteins were measured in cell culture supernatants from treated iBMDMs using ELISAs for IL-1β and CCL5 (Quantikine®, R&D systems).

### siRNA knockdown of SR-BI

iBMDMs were incubated with transfection reagent alone (Lipofectamine® RNAiMAX, 13778150, Thermofisher), scrambled control siRNA (sc-37007, Santa Cruz) or SR-BI siRNA (sc-44753, Santa Cruz) in optiMEM at 100 nM of siRNA for 6h at 37°C. Media was replaced and cells incubated for a total of 48h. iBMDMs were incubated with PBS or discoidal Por-HDL-NPs (10 μg/mL, R4F concentration, 24h) then stimulated with LPS for 16h for inflammatory gene or Por-HDL-NPs for 5h for cellular uptake assessments. Por-HDL-NP uptake was quantified in iBMDM lysates on a Glowmax microplate reader (Promega, excitation λ: 405 nm, emission λ: 660-720 nm), normalized to total protein.

### Western blotting

Western blotting measured nuclear NF-κB p65 and SR-BI. For nuclear NF-κB p65, nuclear protein was isolated using the NE-PER nuclear kit (Thermo Scientific), then proteins separated on 4-12% Bis-Tris Bolt gels using electrophoresis before transfer to nitrocellulose membranes (Thermo Fisher Scientific) by iBlot (Invitrogen). Membranes were incubated with rabbit anti-NF-κB p65 antibody [E379] (1:1000, ab32536, Abcam) followed by goat anti-rabbit HRP-conjugated antibody (1:2000). An anti-rabbit antibody against TATA-binding protein (1:1000, ab63766, Abcam) was the nuclear protein loading control. To validate siRNA SR-BI knockdown, membranes were incubated with anti-SR-BI antibody [EP1556Y] (1:1000, ab52629) followed by goat anti-rabbit HRP-conjugated antibody (1:2000). An anti-alpha tubulin antibody [DM1A] (1:1000, ab7291, Abcam) followed by goat anti-mouse HRP-conjugated antibody were used for protein loading control.

### Animal care

All experiments and procedures were approved by the South Australian Health and Medical Research Institute (SAHMRI) Animal Ethics Committee (AEC application SAM422.19) and conformed to the Australian code for the care and use of animals for scientific purposes. Male C57Bl6/J apolipoprotein (*Apo*)*e*^-/-^ mice bred at SAHMRI were fed high cholesterol diet (HCD) containing 21% fat and 0.15% cholesterol (SF-00219, Semi-Pure Rodent Diet, Specialty Feeds) or standard rodent chow. For early-stage atherosclerosis, 6-week-old mice were fed HCD for a total of 6 weeks. For the unstable plaque model, 12-week-old mice were fed HCD for a total of 13 weeks. Mice were randomized into PBS or discoidal Por-HDL-NPs treatment groups. Blinding was used via random number allocation for histological and flow cytometric analyses.

### Tandem stenosis surgery

After 6 weeks of HCD, mice received tandem stenosis carotid artery ligation surgery as described previously^20^. Briefly, tandem stenoses were made in the right carotid artery with an outer diameter of 150 μm, separated by 3 mm, using 6-0 braided polyester sutures.

After surgery, mice received PBS or discoidal Por-HDL-NPs (40 mg/kg, R4F concentration) via intraperitoneal injection on alternate days for seven weeks until euthanasia.

### 64Cu-Por-HDL-NP γ-counting for biodistribution assessment

*Apoe^-/-^* mice were fed a HCD or chow for 32 weeks. Mice were administered ^64^Cu-labelled 30 mol % Por-HDL-NPs (∼18.5 MBq) and sacrificed 48h post-injection. γ-counts were measured from tissues and calculated as per: 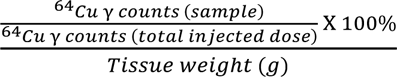

### Positron Emission Tomography (PET)/ Magnetic Resonance Imaging (MRI)

Infusions of ^64^Cu-Por-HDL-NPs (18.5-20 MBq) were administered intravenously into *Apoe^-/-^* mice at week 7 and 11 of HCD feeding or week 12 of HCD feeding for tandem stenosis model. Mice were imaged 6h post-injection with a 5 min scan using the Albira PET (Bruker, MA, USA). PET images were quantified by PMOD v3.509 software (PMOD technologies LLC, Zurich, Switzerland). Volume of interest (VOI) regions were defined using VOI tool. For MRI, mice were imaged with an ICON 1T MRI (Bruker) for anatomical reference to complement PET imaging.

### IVIS fluorescence imaging *ex vivo*

Tissues were harvested 24h after a final intraperitoneal injection of PBS or 5mg/kg of Por-HDL-NP for imaging on the In Vivo Imaging System (IVIS) Spectrum (PerkinElmer, MA, USA) at excitation and emission wavelengths 670 nm and 720 nm, respectively. Epi-fluorescence images were acquired with auto-exposure, F/stop = 2 and smallest binning settings with a field of view of 6.6 - 23 cm. Images were analyzed for fluorescence in regions of interest (ROI) with Aura imaging software (Spectral Instruments Imaging, AZ, USA).

### Histology

Fresh frozen sections of aortic sinus and tandem stenosis Segment I were stained with H&E (plaque area), Masson’s Trichrome (collagen) and Oil red O (lipid). Plaque smooth muscle cells (α-SMA^+^) and macrophages (CD68^+^) were detected by immunofluorescence, and erythrocytes (TER-119^+^) by immunohistochemistry. Sections were imaged under brightfield or fluorescence microscopy and porphyrin-lipid detected with Cy5 filter. Plaque area was calculated from H&E-stained sections. For histological and immunochemical analyses, positive staining was calculated as a percentage of plaque area. See Supplemental Methods for further detail.

### Flow cytometry of aorta and blood

Aortas were digested in Liberase TM (Roche) and passed through a cell strainer (40 µm) then incubated with fluorochrome-conjugated antibodies (Table S3) to detect: macrophages (CD11b^+^F4/80^+^) and M1 (CD86^+^CD206^-^)/M2 (CD86^-^CD206^+^) phenotypes, monocytes (CD11b^+^F4/80^-^), endothelial cells (CD11b^-^ F4/80^-^ CD31^+^) and smooth muscle cells (CD11b^-^F4/80^-^ α-SMA^+^). For blood, taken by tail vein nick in fifth week of treatment in early-stage group, we assessed: neutrophils (CD45^+^ CD11b^+^Gr-1^+^Ly6G^+^), monocytes (CD45^+^CD11b^+^CCR2^+^), activated monocytes (CD45^+^CD11b^+^CCR2^+^Ly6C^hi^). Porphyrin-lipid^+^ cells were detected on the AF647 channel (excitation 640 nm, emission 670/30 nm). Viability dye (AF700) was used for all samples.

Samples were run on a BD LSRFortessa™ X-20 Analyzer and data analyzed with FlowJo™ v10.8.1 software. Fluorescence minus one (FMO) samples were used to set gates with porphyrin-lipid fluorescence gated using PBS controls.

### Statistical analysis

All data expressed as Mean ± SD. Data with groups of more than two were analyzed by one-way ANOVA with post-hoc Tukey’s multiple comparisons. Data with groups of two were analyzed by a two-tailed unpaired t-test, with exception of longitudinal PET activity quantification analyzed using a paired t-test.

## Results

### Por-HDL-NPs and their internalization by macrophages

HDL particles exist in different conformations. When lipid-free apoA-I first acquires lipid is becomes discoidal in shape. Following acquisition of esterified cholesterol into the core, it becomes spherical. These differences affect HDL functionality^11,21^. Accordingly, we synthesized and tested discoidal and spherical-like cholesterol oleate (CO)-loaded Por-HDL-NPs. Their size and shape verified by transmission electron microscopy (TEM, Fig. 1A-B) and absorbance spectra measured (Fig. 1C). Incubation of iBMDMs with discoidal Por-HDL-NPs *in vitro* led to robust uptake. Por-HDL-NPs were visualized within iBMDMs via porphyrin-lipid fluorescence detectable in the cytoplasm of macrophages and absent in the PBS-treated vehicle control (Fig. 1D). Por-HDL-NP uptake was also measurable by flow cytometry (670/30 nm channel, Fig. 1E). iBMDMs incubated with discoidal Por-HDL-NPs exhibited a substantial increase in median fluorescence intensity of the porphyrin-lipid, compared to PBS controls (Fig. 1F, *P*<0.0001).

**Figure 1.**
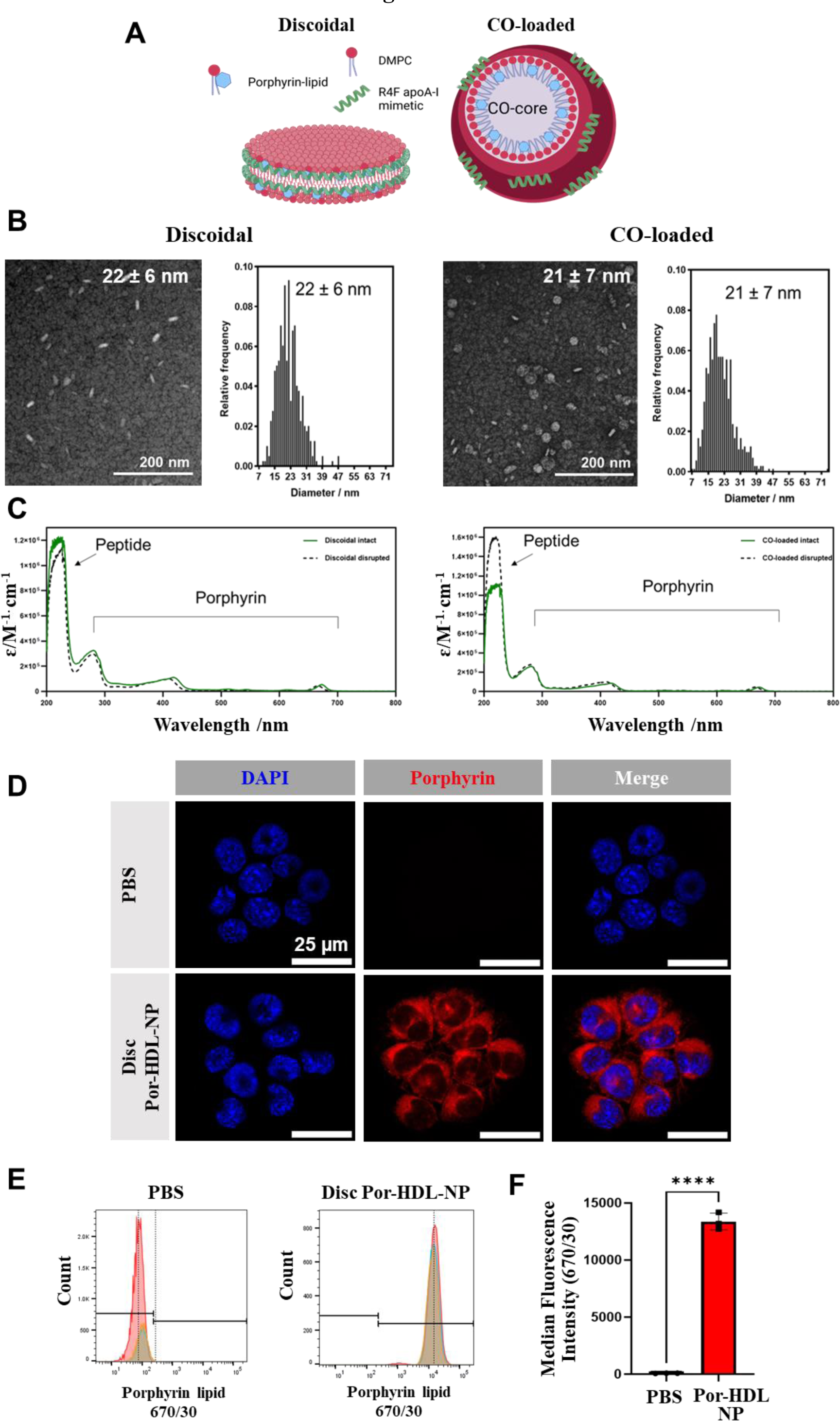
Por-HDL-NPs and their internalization by macrophages *in vitro*. **(A)** Components and structures of discoidal and CO-loaded Por-HDL-NPs. **(B)** Transmission electron microscopy (TEM) of discoidal and CO-loaded Por-HDL-NPs with size distribution, as measured by TEM. **(C)** Absorbance measured at 200-800 nm of either intact (green) or disrupted (dashed) for discoidal and CO-loaded nanoparticles. **(D)** Confocal microscopy images of iBMDMs treated with PBS or discoidal Por-HDL-NPs with porphyrin-lipid detected at 653 nm excitation. **(E)** Flow cytometry of iBMDMs treated with either PBS or discoidal Por-HDL-NPs with porphyrin-lipid fluorescence detected using red laser (640 nm) and fluorescence measured with the 670/30 channel. **(F)** Median fluorescence intensity (MFI) was quantified from flow cytometric analysis of the porphyrin-lipid fluorescence within iBMDMs. Data expressed as Mean ± SD (n=3). \*\*\*\**P*<0.0001 vs PBS control by unpaired two-tailed t-test. Scale bar = 25 µm. DMPC, 1,2-Dimyristoyl-sn-glycero-3-phosphocholine.

### Por-HDL-NPs promote cholesterol efflux and inhibit inflammation in macrophages

Por-HDL-NPs are structural mimetics of HDL, we therefore tested their cholesterol efflux capabilities from iBMDMs. Compared to PBS control, incubation with discoidal and CO-loaded Por-HDL-NPs at 10 μg/mL increased cholesterol efflux (Fig. 2A, Disc: +219%; CO: +189%, *P*<0.0001). Incubation with 25 μg/mL of Por-HDL-NPs led to further stepwise increases in cholesterol efflux compared to PBS control (Fig. 2A, Disc: +415%; CO: +457%, *P*<0.0001). Interestingly, 25 μg/mL of Por-HDL-NPs elicited more cholesterol efflux than 25 μg/mL of rHDL (Fig. 2A, Disc: +49%, *P*<0.001; CO: +61%, *P*<0.0001).

**Figure 2.**
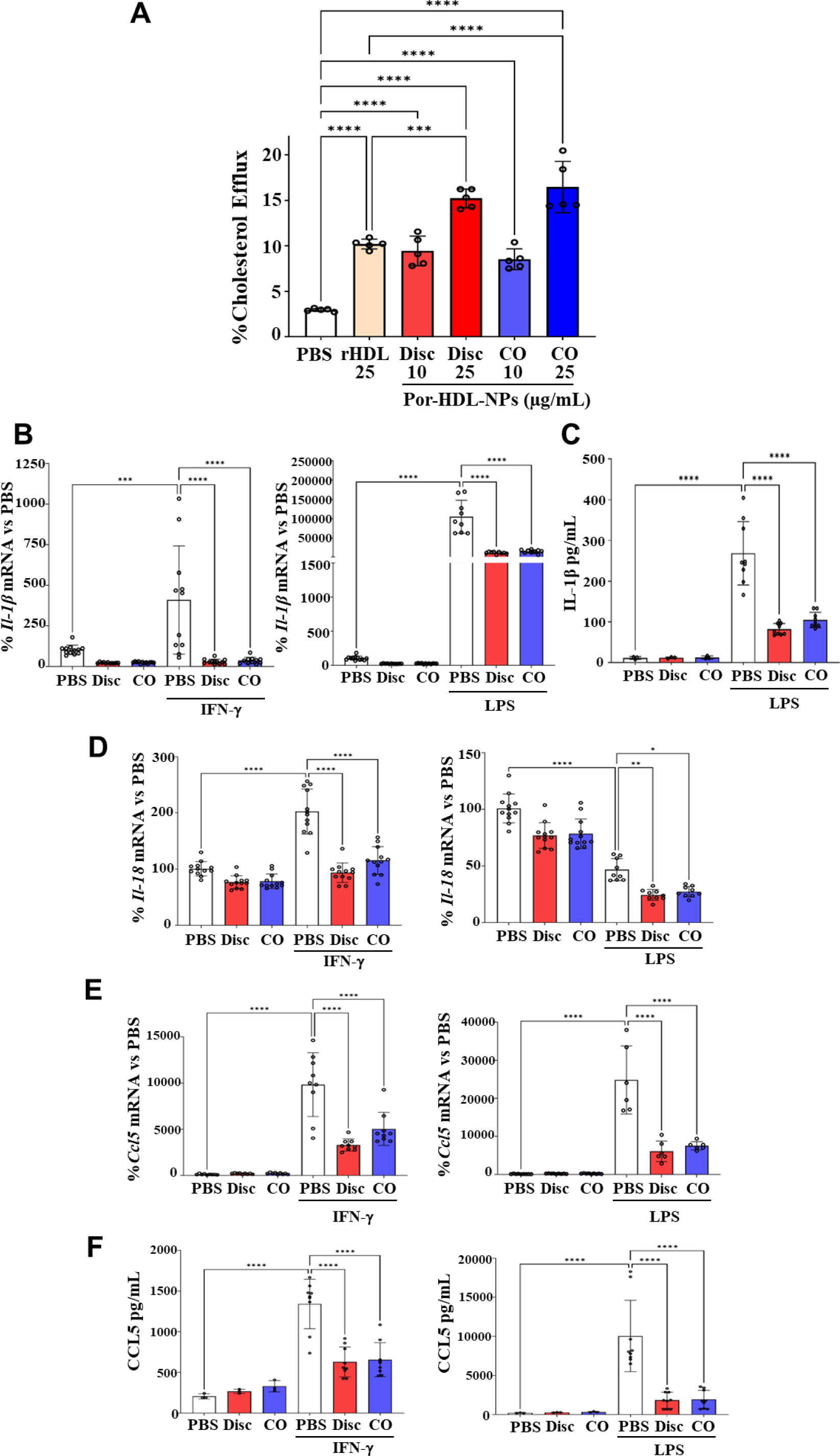
Por-HDL-NPs promote cholesterol efflux and inhibit inflammatory mediators in macrophages *in vitro*. **(A)** Cholesterol efflux measured in [^3^H]-cholesterol-loaded iBMDMs following treatment with either PBS vehicle control, rHDL, discoidal (Disc) or CO-loaded (CO) Por-HDL-NPs (n=5). **(B)** RT-qPCR measurement of *Il-1b* mRNA levels following incubation with discoidal (Disc) or CO-loaded (CO) Por-HDL-NPs and stimulation with IFN-γ (10 ng/mL) or LPS (10 ng/mL). **(C)** ELISA measurement of secreted IL-1β protein in culture media. Following stimulation with IFN-γ and LPS, RT-qPCR assessment of **(D)** *Il-18* and **(E)** *Ccl5* mRNA (n=9-12) and, **(F)** by ELISA, secreted CCL5 protein (n=3-9). Data expressed as Mean ± SD. \**P*<0.05, \*\**P*<0.01, \*\*\**P*<0.001, \*\*\*\**P*<0.0001 by one-way ANOVA with Tukey’s multiple comparisons.

We next determined the anti-inflammatory effects of Por-HDL-NPs in iBMDMs. As expected, IFN-γ and LPS significantly increased *Il-1b* (*P*<0.0001, Fig. 2B). This elevation was attenuated following incubation with discoidal Por-HDL-NPs (IFN-γ: −92%, LPS: −88%, *P*<0.0001) and CO-loaded Por-HDL-NPs (IFN-γ: −91%, LPS: −85%, *P*<0.0001). Consistent with this, secreted IL-1β protein levels were significantly lower in the culture media of iBMDMs incubated with Por-HDL-NPs (Fig. 2C, Disc: −69%; CO: −61%, *P*<0.0001).

iBMDMs preincubated with discoidal and CO-loaded Por-HDL-NPs displayed significantly lower *Il-18* mRNA levels than IFN-γ stimulated controls (Fig. 2D, Disc: −54%; CO: −43%, *P*<0.0001). Although incubation of iBMDMs with LPS caused an unexpected decrease in *Il-18* (Fig. 2D, −54%, *P*<0.001), treatment with both discoidal and CO-loaded Por-HDL-NPs caused an additional decrease in *Il-18* mRNA expression (Fig. 2D, Disc: −48%, *P*<0.01; CO-loaded: 42%, *P*<0.05).

Por-HDL-NPs reduced mRNA levels of chemokine *Ccl5* in iBMDMs for both discoidal (IFN-γ: −66%; LPS: −75%, *P*<0.0001) and CO-loaded (IFN-γ: −49%; LPS: −70%, *P*<0.0001) Por-HDL-NPs, compared to stimulated PBS controls (Fig. 2E). Por-HDL-NPs also decreased secreted CCL5 protein in culture media following stimulation with IFN-γ (Disc: −53%; CO: −51%, *P*<0.0001) and LPS (Disc: −82%; CO: −81%, *P*<0.0001) (Fig. 2F).

### Role of SR-B**I** and cholesterol efflux in the uptake and anti-inflammatory properties of Por-HDL-NPs in macrophages

We next determined whether the uptake and anti-inflammatory properties of Por-HDL-NPs in macrophages were mediated via SR-BI, the target receptor of R4F, *in vitro*.

First, we confirmed that SR-BI mRNA and protein were lower in SR-BI siRNA transfected cells than scrambled (Scr) siRNA controls (Supplemental Fig. S1, mRNA: −59%; protein: −87%, *P*<0.0001). Despite a 37% reduction in Por-HDL-NP fluorescence in SR-BI siRNA iBMDMs, indicating uptake, this did not reach significance (Fig. 3A, *P*=0.07). This suggests non-specific phagocytosis of Por-HDL-NPs by macrophages. Furthermore, significant reductions in *Il-1b* and *Ccl5* mRNA levels were still observed in SR-BI siRNA transfected iBMDMs following treatment with Por-HDL-NPs (Fig. 3B-C), suggesting the anti-inflammatory effects of Por-HDL-NPs are SR-BI-independent.

**Figure 3.**
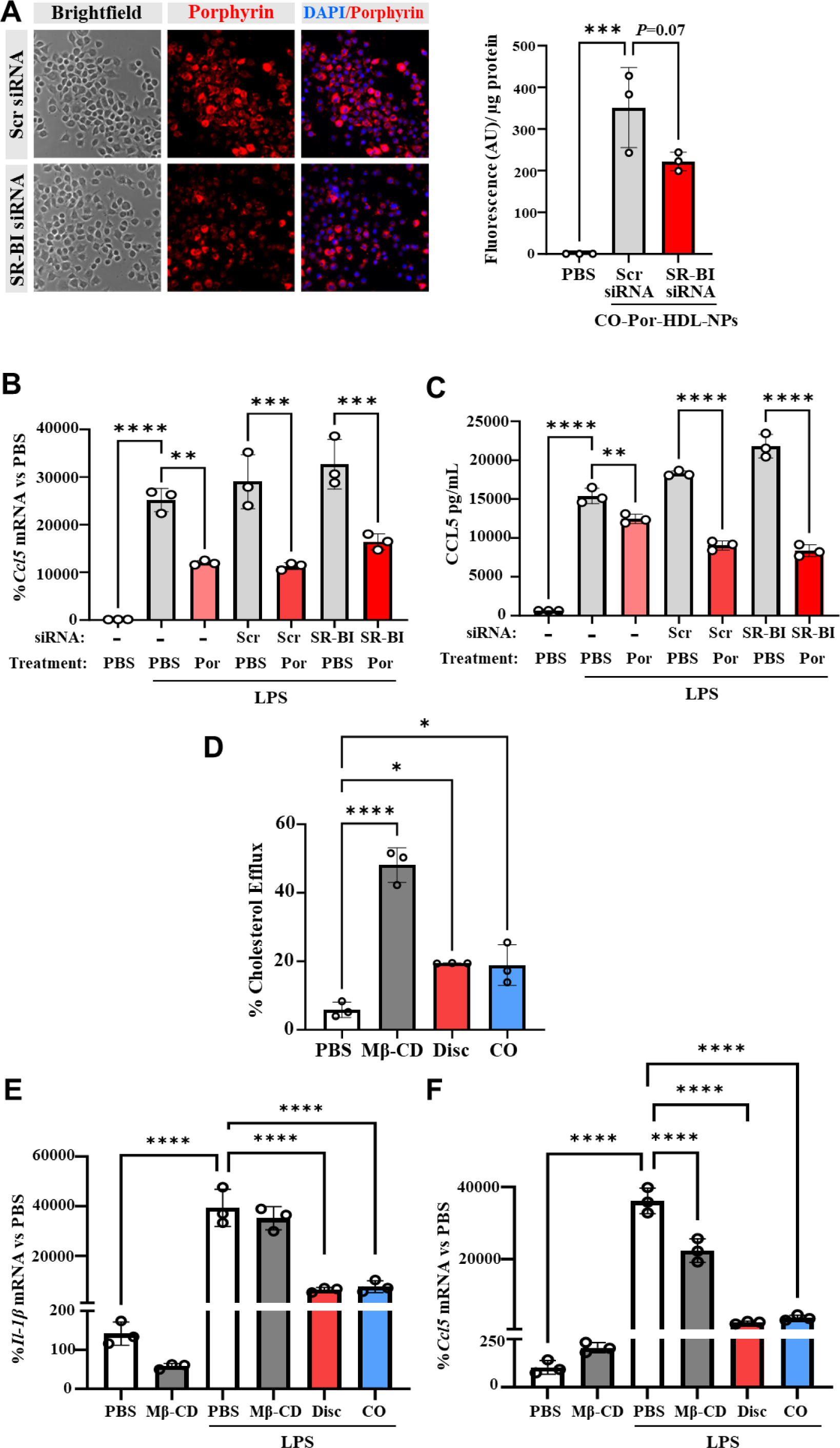
siRNA knockdown of SR-BI and cholesterol efflux have minimal effects on the uptake or anti-inflammatory properties of Por-HDL-NPs. iBMDMs were transfected with SR-BI siRNA (100 nM) to knockdown SR-BI. **(A)** Representative fluorescence microscopy images of Por-HDL-NP uptake following SR-BI siRNA knockdown in iBMDMs with porphyrin fluorescence quantification of uptake in cell lysates (excitation 405nm/ emission 660-720nm, n=3). RT-qPCR measurement of *Ccl5* mRNA **(B)** and ELISA assessment of CCL5 protein **(C)** following incubation with discoidal (Disc) or CO-loaded (CO) Por-HDL-NPs and stimulation with LPS (10 ng/mL) (n=3). **(D)** Cholesterol efflux was measured in iBMDMs following treatment with PBS (control), methyl-β-cyclodextrin (MβCD, 4% v/v), discoidal (Disc) or CO-loaded (CO) Por-HDL-NPs (25 μg/mL). iBMDMs were pre-incubated with PBS, MβCD (4% v/v), discoidal or CO-loaded Por-HDL-NPs (10 μg/mL) then stimulated with LPS (10 ng/mL) before RT-qPCR quantification of the mRNA levels of *Il-1b* **(E)** and *Ccl5* **(F)**. Data expressed as Mean ± SD. \**P*<0.05, \*\**P*<0.01, \*\*\**P*<0.001, \*\*\*\**P*<0.0001 by one-way ANOVA with Tukey’s multiple comparisons.

We next assessed the role of cholesterol efflux in the anti-inflammatory actions of Por-HDL-NPs using passive cholesterol depletion agent methyl-β-cyclodextrin (MβCD). MβCD removed 48% of the loaded [^3^H]-cholesterol, confirming its efflux capabilities (Fig. 3D, 8-fold, vs PBS control, *P*<0.0001). This was greater than the cholesterol efflux induced by discoidal or CO-loaded Por-HDL-NPs. With LPS stimulation, *Il-1b* mRNA levels were not different between the PBS and MβCD treatment groups (Fig. 3E), yet Por-HDL-NPs caused significant reductions in *Il-1b* mRNA (Fig. 3E, Disc: −84%; CO: −81%, *P*<0.0001). However, MβCD significantly decreased *Ccl5* mRNA levels (Fig. 3F, −38%, p<0.0001), compared to LPS stimulated controls, but Por-HDL-NPs caused a greater reduction in *Ccl5* mRNA levels (Fig. 3F, Disc: −89%; CO: −84%, *P*<0.0001). These findings suggest inhibition of *Ccl5* but not *Il-1b* by Por-HDL-NPs may be a result of cholesterol efflux, but other mechanisms are also operating.

### Por-HDL-NPs suppress NLRP3 inflammasome components and p65-Nf-κB

Por-HDL-NPs supressed NLRP3 inflammasome-associated cytokines IL-1β and IL-18. We next measured changes in key components of this pathway. Incubation of iBMDMs with Por-HDL-NPs significantly lowered *Nlrp3* mRNA levels (Fig. 4A, Disc: −69%, p<0001; CO: −62%, p<0.001) compared to IFN-γ stimulated controls. No changes in *Nlrp3* mRNA levels were observed with Por-HDL-NP incubations following LPS stimulation (Fig. 4B). Discoidal Por-HDL-NPs significantly reduced *Asc* mRNA levels in response to IFN-γ stimulation (Fig. 4C, −36%, *P*<0.01). In iBMDMs stimulated with LPS, discoidal and CO-loaded Por-HDL-NPs reduced *Asc* mRNA levels as compared to the PBS non-stimulated controls (Fig. 4D, Disc: −56%, p<0001; CO: −55%, p<0.001). Por-HDL-NPs caused no significant changes in *Asc* mRNA levels when compared to LPS-stimulated controls (Fig. 4D).

**Figure 4.**
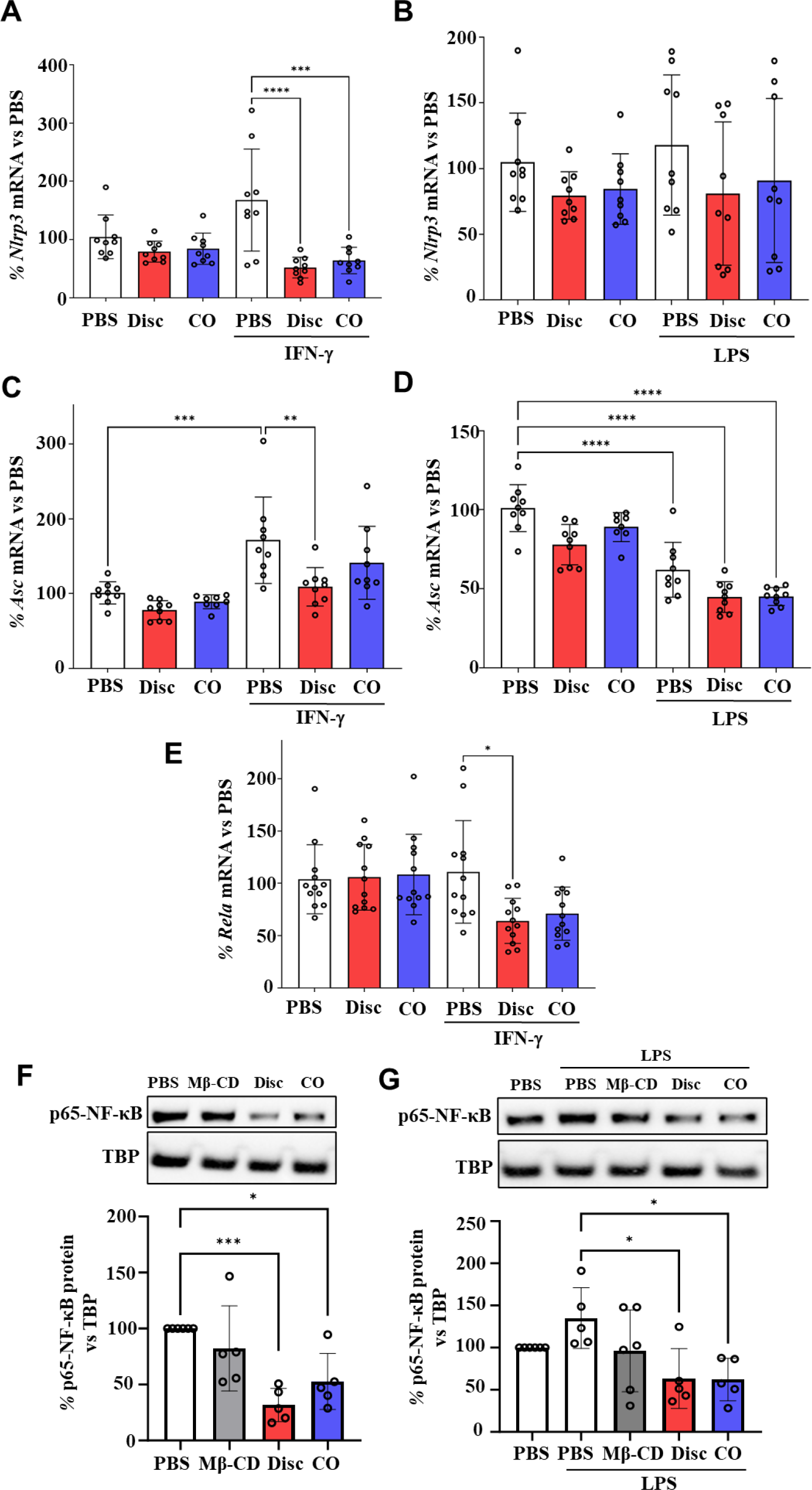
Por-HDL-NP suppress components of the NLRP3 inflammasome and reduce p65-NFκB activation. RT-qPCR measurements of **(A-B)** *Nlrp3*, **(C-D)** *Asc* and **(E)** *Rela* mRNA levels following incubation with discoidal (Disc) or CO-loaded (CO) Por-HDL-NPs and stimulation with IFN-γ (10 ng/mL) or LPS (10 ng/mL) (n=9-12). The nuclear fractions of **(F)** unstimulated and **(G)** LPS-stimulated iBMDMs were isolated and nuclear p65-NFκB protein levels (∼65 kDa) were assessed by Western blotting (n=5-6). TATA binding protein (TBP, 37 kDa) was used as the nuclear protein loading control. Data expressed as Mean ± SD. \**P*<0.05, \*\*\**P*<0.001, \*\*\*\**P*<0.0001 by one-way ANOVA with Tukey’s multiple comparisons.

We next measured mRNA levels of *Rela*, the gene that transcribes the p65 active subunit of NFκB. Stimulation with IFN-γ did not increase *Rela* mRNA levels, but in those cells incubated with discoidal Por-HDL-NPs we observed a significant decrease in *Rela* (Fig. 4E, −42%, *P*<0.05), compared to the stimulated PBS control. When NFκB is activated, the p65 subunit translocates to the nucleus to upregulate inflammatory gene expression. We next determined changes in p65-NFκB protein in the nuclear fraction of iBMDMs. In non-stimulated iBMDMs, incubation with Por HDL-NPs caused a reduction in nuclear p65-NFκB protein levels compared to the PBS control (Fig. 4F, Disc: −68%, *P*<0.0001; CO: −47%, *P*<0.05). There was no change in nuclear p65-NFκB protein in response to MβCD in non-stimulated iBMDMs. In LPS-stimulated iBMDMs, both discoidal and CO-loaded Por HDL-NPs reduced nuclear p65-NFκB protein levels (Fig. 4G, Disc: −53%, *P*<0.0001; CO: −54%, *P*<0.05), compared to LPS control cells. This effect was independent of cholesterol efflux as incubation with MβCD did not cause a significant reduction in nuclear p65 protein (Fig. 4G, −29%, *P*=0.33).

Overall, Por-HDL-NPs suppress the activation of the inflammatory transcription factor p65-NFκB and components of the NLRP3 inflammasome, all of which play important roles in atherosclerosis.

### Detection of Por-HDL-NPs in murine plaques using multimodal imaging

To determine the biodistribution of Por-HDL-NPs *in vivo*, we infused ^64^Cu-labelled Por-HDL-NPs into *Apoe^-/-^* mice fed a chow or high cholesterol diet (HCD) for 32 weeks. *Apoe^-/-^* mice fed HCD will have significantly more atherosclerosis than chow-fed mice^22^. For chow and HCD mice, the highest uptake of ^64^Cu-labelled Por-HDL-NPs was in the liver (Fig. 5A). Uptake of ^64^Cu-Por-HDL-NPs was significantly higher in the heart of HCD-fed mice, compared to chow-fed mice (Fig. 5B, +15%, *P*<0.05). There was a non-significant trend for higher uptake of ^64^Cu-Por-HDL-NPs in the aortas of mice fed HCD compared to chow (Fig. 5B, +51%, *P*=0.0623). We found significantly higher uptake of ^64^Cu-Por-HDL-NPs in blood (Fig. 5B, +34%, *P*<0.05) and gut (Fig. 5B, +39%, *P*<0.05) in HCD-fed mice compared to chow controls. Overall, this provides evidence for increased localization of Por-HDL-NPs to regions where plaque accumulates (heart/ aorta) in HCD-fed *Apoe*^-/-^ mice and in circulating cells that contribute to plaque development.

**Figure 5.**
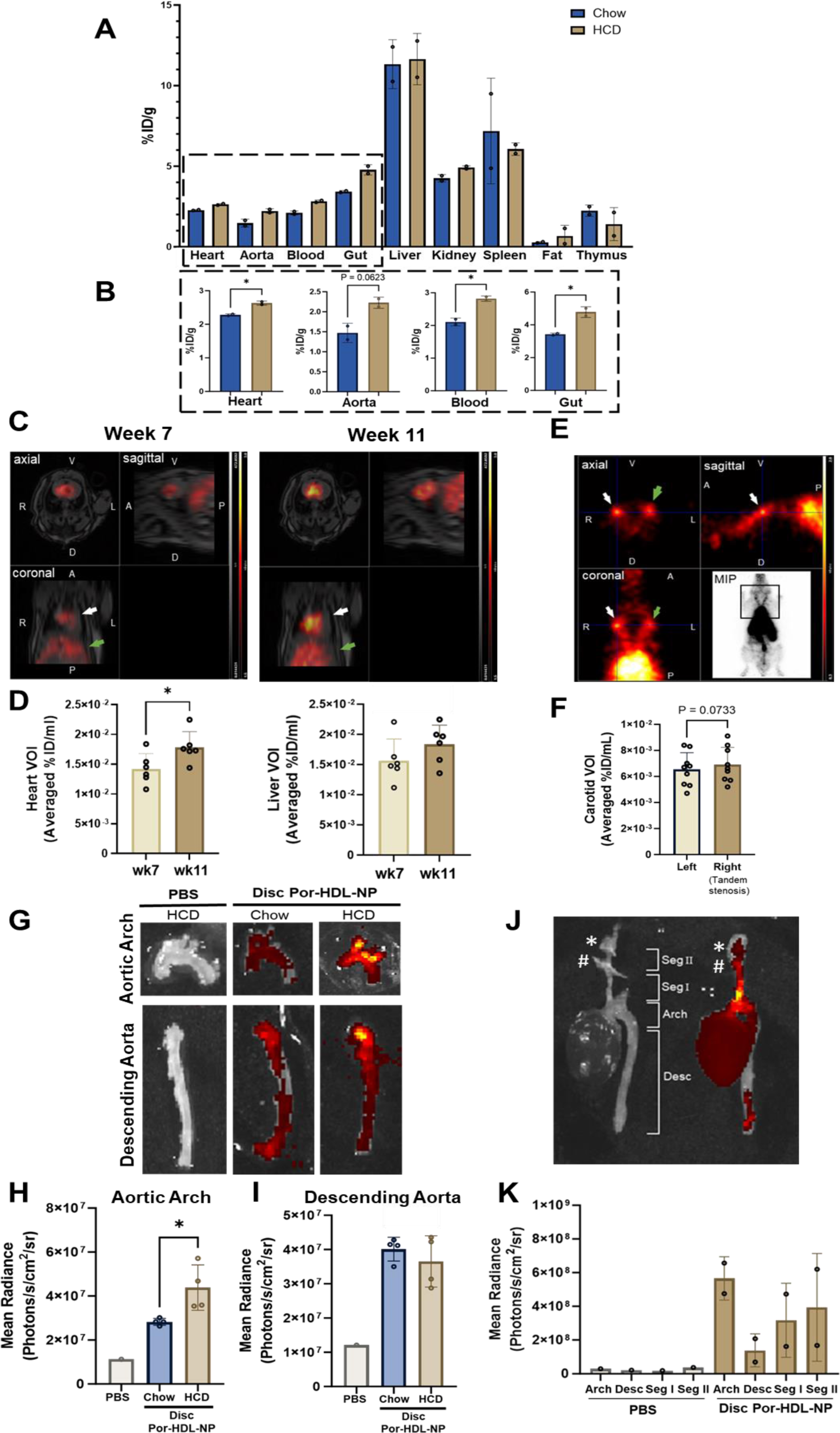
Multimodal imaging of Por-HDL-NPs in atherosclerosis. **(A)** γ-counting of ^64^Cu-Por-HDL-NPs in organs from *Apoe^-/-^* mice fed either a chow or HCD for 32 weeks (n=2/group). **(B)** Comparisons of γ-counts in heart, aorta, blood and gut between chow and HCD-fed mice. **(C)** Merged PET/MRI images in the axial, sagittal and coronal planes in HCD fed *Apoe^-/-^* mice after week 7 and week 11 of HCD, and 6h after intravenous administration of ^64^Cu-Por-HDL-NPs. White arrows show signal in the heart region and green arrows show signal in part of the liver region. **(D)** VOI measurements of PET signal in heart and liver regions at week 7 and week 11 of HCD (n=6). **(E)** Representative PET images in the axial, sagittal and coronal planes at 6 weeks post-surgery (week 12 of HCD) of the left and right carotid in *Apoe-/-* mice with a tandem stenosis in the right common carotid artery, 6 h post-intravenous injection of ^64^Cu-Por-HDL-NPs. White and green arrows indicate the right and left common carotid artery respectively. **(F)** Quantification of the PET activity in the left and right carotid VOI in *Apoe^-/-^* tandem stenosis mice 6h post-injection of ^64^Cu-Por-HDL-NPs. Data expressed as mean ± SD (n=9). *P*>0.05 vs left carotid by unpaired two-tailed t-test. **(G)** Representative images of IVIS fluorescence images captured side-by-side in one image frame of the excised aortic arch and descending aorta of PBS or Por-HDL-NP infused mice fed either chow or HCD for 8 weeks. Quantification of the fluorescence signal from images of excised aortic arch **(H)** and descending aorta **(I)** expressed as mean radiance (Photons/s/cm2/sr). n=1-4 mice/group (PBS, n=1; Por-HDL-NP Chow, n=1; Por-HDL-NP HCD, n=4). Fluorescence imaging of Por-HDL-NP Chow and HCD aortic arches and descending aortas were captured side-by-side in a single image with the IVIS camera. (**J**) IVIS fluorescence images of the vascular tree of *Apoe^-/-^* mice with tandem stenoses in the right common carotid artery with distal (*) and proximal (#) sutures indicated, and tandem stenosis Segments I & II (Seg I & II) shown. (**K**) Quantified PET activity throughout the vascular tree with tandem stenoses comparing PBS and Por-HDL-NP (n=2). Data expressed as Mean ± SD.\**P<*0.05 by unpaired two-tailed t-test. V, Ventral; D, Dorsal; R, Right; L, Left; A, Anterior; P, Posterior. ID, Injected dose. VOI, Volume of interest. MIP, Maximal Image Projection.

We next sought to determine whether ^64^Cu-labelled Por-HDL-NPs track to areas of plaque and whether they can be used to assess changes in plaque size longitudinally over time (Fig. 5C-D). PET/MRI imaging of *Apoe^-/-^* mice fed a HCD demonstrated a quantifiable increase in PET signal in the heart VOI from week 7 to week 11 of HCD feeding (Fig. 5D, Week 7, 1.42×10^-2^ %ID/mL *vs*. Week 11, 1.741×10^-2^ %ID/mL, *P*<0.05). There was no change in PET signal in the liver VOI (Fig. 5D, Week 7, 1.567×10^-2^ %ID/mL; Week 11, 1.864×10^-2^ %ID/mL). We next examined whether Por-HDL-NPs localized to areas of plaque instability in mice subjected to tandem stenosis surgery. ^64^Cu-Por-HDL-NP activity was detected in the left and right carotid regions using PET imaging (Fig 5E). When comparing left and right carotid regions, there was a non-significant increase in ^64^Cu-Por-HDL-NP activity in the right carotid with tandem stenosis-induced plaque (Fig 5F, Left, 6.533×10^-3^ %ID/mL; Right, 6.9×10^-3^ %ID/mL; P=0.0733), compared to the left carotid.

We next explored the fluorescence imaging capabilities of Por-HDL-NPs and ability to track to plaque in HCD-fed *Apoe^-/-^* mice. Using IVIS fluorescence imaging, excised aortic arches and descending aortas were compared following infusion of PBS or Por-HDL-NPs (Fig. 5G). HCD-fed mice elicited a stronger fluorescence signal in the arch region, where plaque develops, than chow-fed Por-HDL-NP control (Fig. 5H, +55%, *P*<0.05). In contrast, there were no changes in fluorescence detected between HCD and chow-fed controls in the descending aorta (Fig. 5I), a site with little plaque after 8 weeks HCD. This demonstrates that the increased fluorescence signal in the aortic arch is specific to Por-HDL-NP uptake into atherosclerotic plaque and not ubiquitous across the aorta. Por-HDL-NP fluorescence was also quantified in the liver, kidney, spleen and lung (Supplemental Fig. S2A). Quantification of the fluorescence signal revealed that Por-HDL-NP fluorescence was greatest in the liver of HCD fed mice compared to chow-fed controls (Fig. S2B, 2-fold, *P*<0.0001). The fluorescence signal from the livers of HCD-fed mice was significantly higher than the other organs including the kidney, spleen and lungs (Fig. S2B, ∼10-fold, *P*<0.0001 for all). In animals with plaque instability induced by tandem stenoses in the carotid, IVIS fluorescence imaging was also conducted *ex vivo* on the vascular tree (Fig. 5J). Fluorescence was quantified from different regions including aortic arch, descending aorta, and tandem stenosis segments I and II, with fluorescence clearly detected in the Por-HDL-NP but not PBS group (Fig. 5K). Fluorescence signal was the highest in the aortic arch, compared to the descending aorta and segment I or II, suggesting that Por-HDL-NPs indiscriminately localize to areas of both stable and unstable plaque.

Next, using fluorescence microscopy we showed that Por-HDL-NP uptake localizes to areas of stable (aortic sinus) and unstable (tandem stenosis Segment I) plaque. In mid-stage stable plaques from the aortic sinus, porphyrin-lipid fluorescence was localized within the lesion, particularly in the shoulder and luminal-facing regions (Fig. 6A). We found porphyrin-lipid fluorescence primarily co-localized to regions of CD68^+^ macrophages in the plaque (Fig. 6A). In very early-stage stable plaque in the aortic sinus, porphyrin-lipid fluorescence was present throughout the majority of the plaque in Por-HDL-NP infused and not PBS-treated mice (Fig. 6B). Moreover, porphyrin-lipid fluorescence was present in plaque of segment I from the carotid artery of the tandem stenosis model in the Por-HDL-NP treated mice and was co-localized with CD68^+^ macrophages (Fig. 6C).

**Figure 6.**
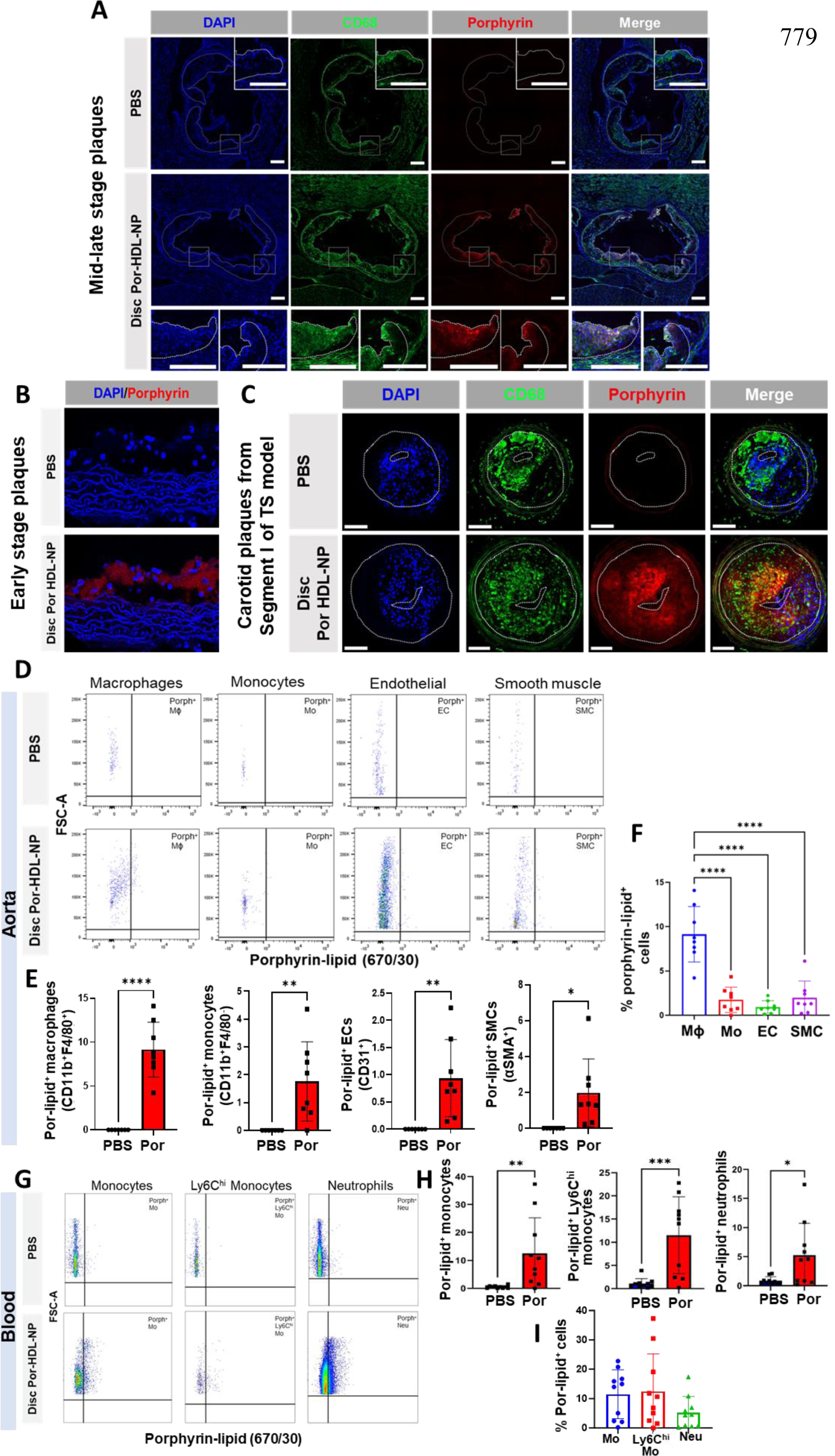
Por-HDL-NP fluorescence detected in atherosclerotic plaques, aortic macrophages and circulating cells *in vivo*. **(A)** After 6 weeks of HCD, *Apoe*^-/-^ mice received tandem stenosis via carotid artery ligation surgery. Mice then received infusions of either PBS or Por-HDL-NP on alternate days for a further 7 weeks. Immunofluorescence was used in fresh frozen mid-late-stage aortic sinus stable plaque sections to identify CD68^+^ macrophages (FITC, green) and nuclei (DAPI, blue). Fluorescence microscopy visualized porphyrin-lipid within aortic sinus sections (Cy5-filter, red) and its colocalization with CD68^+^ macrophages (white). Scale bar = 200 µm. **(B)** *Apoe*^-/-^ mice were fed HCD for 6 weeks and during that time received PBS or Por-HDL-NP infusions three times weekly. Porphyrin lipid fluorescence was identified in early-stage plaques by fluorescence microscopy. **(C)** The presence of porphyrin lipid (red), CD68^+^ macrophages (green) and their colocalization (yellow) was assessed in carotid artery plaque from the tandem stenosis model. Scale bar = 100 µm. **(D)** The descending thoracic aortas from were subjected to flow cytometry analysis for porphyrin lipid fluorescence in aortic: **(E)** CD11b^+^F4/80^+^ macrophages (MΦ), CD11b^+^F4/80^-^ monocytes (Mo), CD31^+^ endothelial cells (ECs) and αSMA^+^ smooth muscle cells (SMCs) (n= 6-8). **(F)** Comparison of porphyrin lipid uptake into aortic cell types. **(G)** Circulating cells in blood were assessed by flow cytometry for porphyrin lipid fluorescence in: **(H)** monocytes, Ly6C^hi^ activated monocytes and Gr-1^+^Ly6G^+^ neutrophils. **(I)** Comparison of porphyrin-lipid uptake into circulating cells (n=8). Data expressed as Mean ± SD. \**P*<0.05, \*\**P*<0.01, \*\*\**P*<0.001, \*\*\*\**P*<0.0001 by unpaired two-tailed t-test or one-way ANOVA with Tukey’s multiple comparisons. Por: Por-HDL-NP.

Using flow cytometry, we detected porphyrin-lipid positive macrophages, monocytes, endothelial cells and smooth muscle cells in digested aortas excised from *Apoe*^-/-^ mice infused with Por-HDL-NPs, which were not present in PBS controls (Fig. 6D-E). In *Apoe^-/-^* mice infused with Por-HDL-NPs, the highest proportion of porphyrin-lipid positive cells were in the macrophage population (9%), with comparatively small amounts in the monocyte (1.8%), endothelial (0.09%) and smooth muscle (2%) cell populations (Fig. 6F, *P*<0.0001). In blood, porphyrin-lipid was detected in circulating monocytes (Fig. 6G-H, 11.5%) and activated monocytes (Fig. 6G-H, 12.5%) and neutrophils (Fig. 6G-H, 5.2%). There were, however, no significant differences in Por-HDL-NP uptake between these circulating cell populations (Fig 6I).

### Por-HDL-NPs exhibit anti-atherosclerotic properties *in vivo*

We next tested the therapeutic effects of Por-HDL-NPs *in vivo* in early-stage plaque and unstable atherosclerosis models. We chose discoidal Por-HDL-NPs due to their minimal *in vitro* differences to CO-loaded spherical-like Por-HDL-NPs. For the early plaque model, in 4-week-old *Apoe*^-/-^ mice, after 6 weeks of treatment, analysis of plaque in the aortic sinus revealed that Por-HDL-NP infused mice had significantly smaller plaques (Fig. 7A, −23%, *P*<0.05) than PBS controls. There were no changes in CD68^+^ macrophage and α-SMA^+^ SMC content in early-stage plaques between treatment groups (Supplemental Fig. S3A-B). In the second tandem stenosis unstable plaque model, *Apoe*^-/-^ mice were treated for 7 weeks following surgery. H&E-stained sections were analyzed at three regions spanning tandem stenosis Segment I^20^ (Fig. 7B). At the mid-point region of Segment I, there was a significant decrease in plaque area in the Por-HDL-NP group, compared to PBS control (Fig. 7D, −52%, *P*<0.05). No differences were observed in the outer regions of Segment I (Fig. 7C&E). There were no changes in CD68^+^ macrophage, Oil Red O lipid, α-SMA^+^ SMC or TER-119^+^ erythrocyte plaque content between groups (Supplemental Fig. S3C-G), with a significant decrease in plaque collagen content in Por-HDL-NP treated mice (Supplemental Fig. S3H).

**Figure 7.**
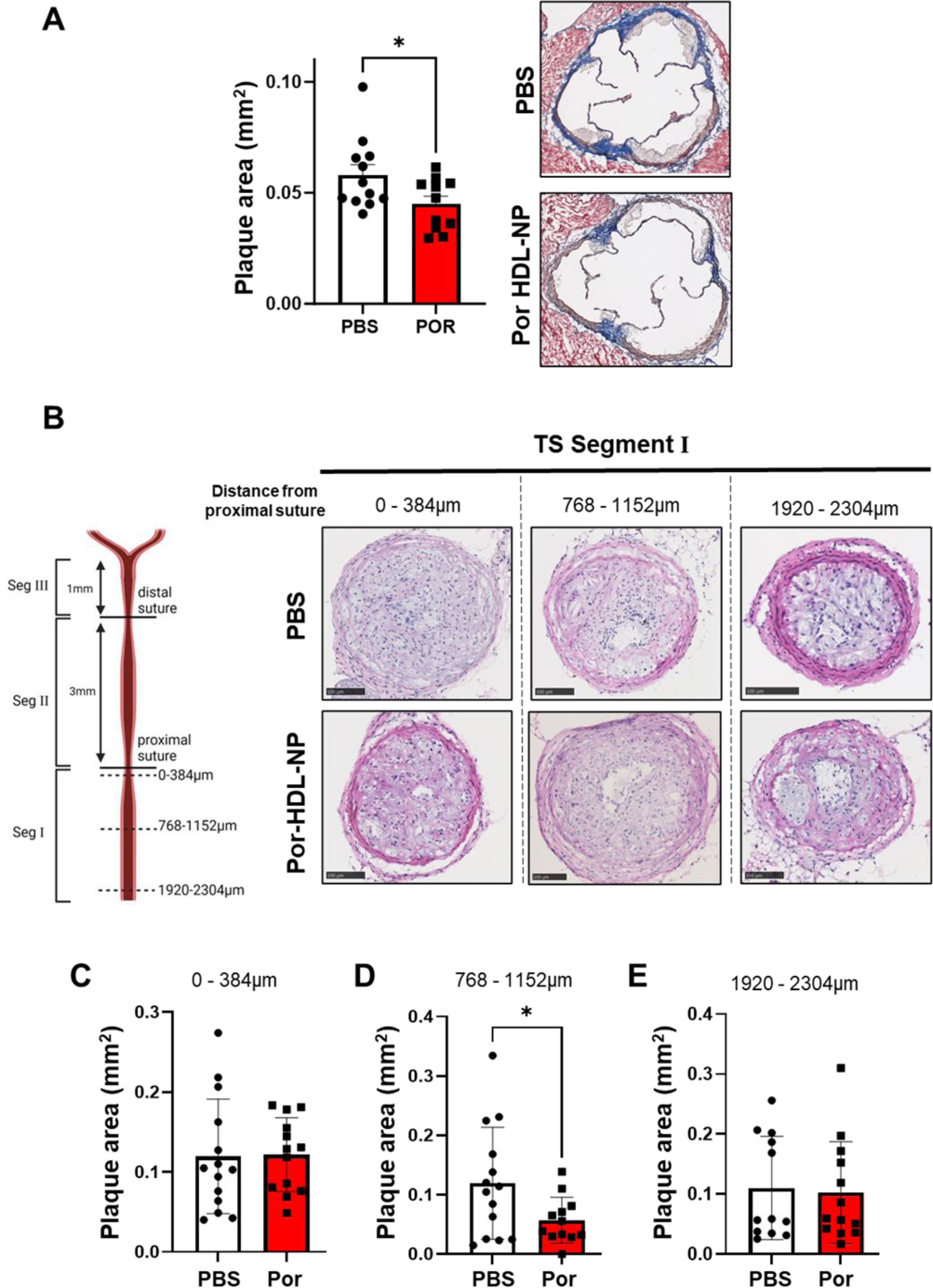
Por-HDL-NPs reduce stable and unstable plaque area. **(A)** Six-week-old *Apoe*^-/-^ mice were fed HCD for 6 weeks to develop early-stage plaque in the aortic sinus and received PBS or Por-HDL-NP infusions three times weekly. Three sections spanning the aortic sinus were stained with Masson’s Trichrome. Plaque area was calculated and averaged across the sections (mm^2^, n=12/group). Right: representative aortic sinus sections stained with Masson’s Trichrome. **(B)** After 6 weeks of HCD, *Apoe*^-/-^ mice received tandem stenosis carotid artery ligation surgery and then infused with PBS or Por-HDL-NPs on alternate days for a further 7 weeks. **Left:** diagram of the sectioned areas of tandem stenosis Segment I in the right common carotid with three regions analyzed for area measurements at the beginning, middle and end of the segment corresponding to 0-384 μm, 768-1152 μm, and 1920-2304 μm from the proximal suture respectively. **Right:** representative images of H&E-stained tandem stenosis Segment I sections within the three regions. Scale bar = 100 µm. Plaque area in tandem stenosis segment I quantified in regions: **(C)** 0-384 μm, **(D)** 768-1152 μm and **(E)** 1920-2304 μm (n=12-14/group). Data expressed as mean ± SD. \**P*<0.05 by unpaired two-tailed t-test. TS, Tandem Stenosis. Por: Por-HDL-NP. Seg: Segment.

We next examined changes in circulating and aortic cells by flow cytometry. In blood, we found a significant reduction in circulating monocytes in mice treated with Por-HDL-NPs (Fig. 8A, −32%, *P*<0.05). However, no changes were observed in activated monocytes (Fig. 8B) or neutrophils (Fig. 8C). Analyses of aortas from treated mice found a significant reduction in monocyte content (Fig. 8D, −81%, *P*<0.05) in mice infused with Por-HDL-NP, when compared to PBS controls. There were non-significant reductions in aortic macrophages (Fig. 8E, −49%, *P*=0.1080) and M1 (Fig. 8F, −73%, *P*=0.051) and M2 (Fig. 8G, −77%, *P*=0.08) macrophage phenotypes, and TREM2^+^ foam cells (Fig. 8H, −50%, *P*=0.13) in Por-HDL-NP treated mice. No changes were observed in aortic endothelial and smooth muscle cells between groups (Fig. 8I-J).

**Figure 8.**
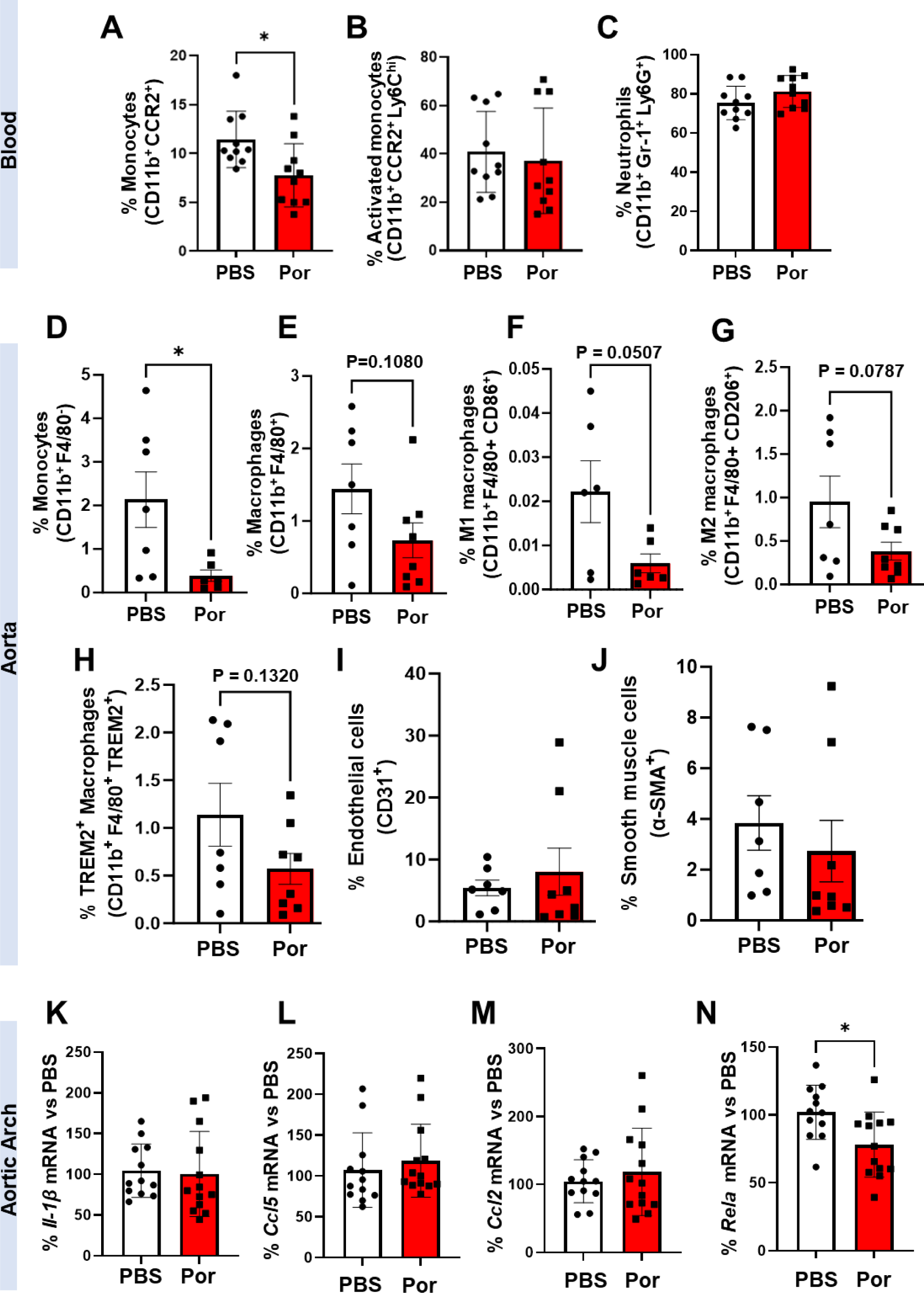
Por-HDL-NPs reduce circulating monocyte and aortic macrophage numbers and inhibit aortic *Rela* expression *in vivo*. Circulating **(A)** monocytes, **(B)** activated monocytes and **(C)** neutrophils were measured and quantified in blood using flow cytometry in mice fed a HCD for 6 weeks (n=10/group). Aortic **(D)** monocytes, **(E)** macrophages, **(F)** M1 macrophages, **(G)** M2 macrophages, **(H)** TREM2^+^ foam cell macrophages, **(I)** endothelial cells and **(J)** and smooth muscle cells were measured and quantified in the descending aorta using flow cytometry in *Apoe-/-* mice fed a HCD for 13 weeks (n=7-8/group). qPCR of RNA extracted from the aortic arch of mice fed a HCD for 13 weeks to determine mRNA levels of: **(K)** *Il-1b,* **(L)** *Ccl5,* **(M)** and (N) *Rela.* Data expressed as mean ± SD. \**P*<0.05 by unpaired two-tailed t-test. POR: Por-HDL-NP.

Whilst no differences were observed in the mRNA levels of inflammatory genes *Il-1b*, *Ccl5* or *Ccl2* in the aortic arches between groups (Fig 8K-M), there was a significant reduction in aortic arch *Rela* mRNA levels in Por-HDL-NP treated mice (Fig. 8N, −26%, *P*<0.05).

In both the early plaque and unstable plaque models, there were no changes in plasma total cholesterol, LDL cholesterol or HDL cholesterol between treatment groups (Supplemental Tables S4 and S5). In the early-stage plaque model, there was a modest (<10%) yet significant increase in body weight in Por-HDL-NP treated mice (Supplemental Table S4, +6.0%, *P*<0.05). Conversely, in the unstable plaque model, there was a significant reduction in body weight (Table S5, −8.6%, *P*<0.05) and plasma triglyceride concentrations (Supplemental Table S5, −22.9%, *P*<0.01) in Por-HDL-NP treated mice.

## Discussion

Nanotechnologies have emerged as highly promising novel CVD treatment and diagnostic strategies with macrophages presenting as desirable cellular targets^5^. A significant advantage of nanoparticles is they can be synthesized to combine therapeutic and diagnostic components into a single theranostic nanoscale platform^6^. This study tested Por-HDL-NPs for their ability to detect and suppress atherosclerosis. We report the following important findings: 1) Por-HDL-NPs are internalized by macrophages, which can be visualized by fluorescence microscopy and detected by flow cytometry; 2) Por-HDL-NPs exhibit anti-atherosclerotic effects *in vitro* in macrophages and promote cholesterol efflux and inhibit inflammation; 3) Por-HDL-NPs localize within plaques *in vivo* and can be detected using PET and fluorescence imaging; and 4) Por-HDL-NP uptake in plaque is primarily by macrophages. 5) Infusions of Por-HDL-NPs reduced plaque size in early-stage stable and tandem stenosis unstable models of atherosclerosis, and decreased circulating and aortic monocytes. Our findings demonstrate the theranostic potential of Por-HDL-NPs for atherosclerosis.

Por-HDL-NPs contain the apoA-I mimetic peptide R4F, which enables interaction with the SR-BI receptor^16^. SR-BI is expressed and functionally utilized by macrophages to facilitate both efflux and influx of cholesterol, and mediates activation of downstream signaling pathways^18^. Similarly, other R4F nanoparticles interact with SR-BI in Chinese hamster ovary cells^16^. Here we observed Por-HDL-NP uptake by macrophages (iBMDMs), consistent with other HDL-mimetic nanoparticles containing full-length apoA-I or R4F^23–25^. To our knowledge, this is the first time R4F-containing Por-HDL-NPs have been shown to be internalized by macrophages, visualized using the fluorescent properties of porphyrin-lipid.

Using PET and fluorescence imaging modalities we found ^64^Cu-Por-HDL-NPs tracked to atherosclerotic plaque *in vivo*. We also found the ^64^Cu-Por-HDL-NP signal was sensitive enough to detect measurable increases in plaque in the heart region. In longitudinal measures on *Apoe*^-/-^ mice, serial PET imaging showed an increase in PET signal in the heart region from 7 to 11 weeks on HCD. A unique feature of the Por-HDL-NP design is that the integrated porphyrin-lipid does not require the conjugation of additional fluorophores^14,15,26^. Using IVIS imaging, we could therefore also detect higher porphyrin-lipid fluorescence signal in the aortic arch of HCD-fed *Apoe^-/-^* mice, compared to chow fed controls with no plaque. This indicates specific uptake in a known region of plaque deposition^27^, supporting the PET observations. Consistent with this, we observed porphyrin-lipid fluorescence in tissue sections of stable plaque and unstable plaque, co-localized with CD68^+^ macrophages, by fluorescence microscopy. The plaque targeting capabilities of Por-HDL-NPs were further demonstrated in biodistribution studies that showed higher activity of ^64^Cu-Por-HDL-NPs in the heart and aorta of HCD-fed mice with established plaque than chow-fed mice with little-to-no plaque. Flow cytometry revealed porphyrin-lipid uptake was highest in aortic macrophages and was detected in circulating monocytes. Circulating monocytes infiltrate and differentiate into macrophages within atherosclerotic lesions^28^, and perhaps porphyrin-lipid fluorescence is retained in this process. In summary, Por-HDL-NPs are taken up preferentially by aortic macrophages and are detected in atherosclerotic plaque, targeting the region of interest.

HDL exhibit anti-inflammatory effects in macrophages via interaction with SR-BI^29^. We found that Por-HDL-NPs also inhibited macrophage expression of pro-inflammatory cytokines IL-1β and IL-18, and the chemokine CCL5. IL-1β and IL-18 have a key role in atherogenesis^30–32^ and their release can be attributed to NLRP3 inflammasome activation^33,34^. Por-HDL-NPs inhibited the expression of NLRP3 inflammasome components, which may in part explain the reduction in IL-1β and IL-18. Interestingly, knockdown of SR-BI did not affect the anti-inflammatory actions of Por-HDL-NPs, despite the reported importance of SR-BI in mediating the anti-inflammatory actions of HDL^29^. Using the passive cholesterol efflux agent, methyl-β-cyclodextrin (MβCD), we found that whilst cholesterol efflux had no effect on the inhibitory effects of Por-HDL-NPs on *Il-1b*, MβCD caused a significant reduction in *Ccl5*. This suggests that cholesterol efflux has some involvement in mediating the inhibitory effects of Por-HDL-NPs on *Ccl5*, but only part, with Por-HDL-NPs causing a more substantial reduction in *Ccl5* than MβCD.

We did find that Por-HDL-NPs suppressed p65-NF-κB activation in iBMDMs and *Rela* in aortic arches *in vivo*. Using MβCD, the inhibitory effect on NF-κB was also independent of cholesterol efflux. NF-κB is the pivotal transcription factor that controls the expression of inflammatory cytokines and chemokines, and also primes NLRP3 inflammasome assembly^34^. The suppression of p65-NF-κB activation by Por-HDL-NPs supports their inhibition of IL-1β, *Il-18* and CCL5, as well as the *Nlrp3* and *Asc*, NLRP3 inflammasome components. Knockdown of SR-BI caused a non-significant reduction (37%) in Por-HDL-NP uptake into iBMDMs. We suggest the pathway by which Por-HDL-NPs inhibit NF-κB is primarily via phagocytotic macrophage entry rather than via SR-BI. Overall, *in vitro* Por-HDL-NPs exhibit anti-inflammatory properties in macrophages through inhibition of NF-κB. This is predominantly independent of interaction with SR-BI or cholesterol efflux.

We found Por-HDL-NPs were highly efficient cholesterol efflux acceptors. By reducing foam cell formation, cholesterol efflux is associated with lower incidence of CAD^35,36^. Taken together, the anti-inflammatory and cholesterol efflux properties of Por-HDL-NPs present them as anti-atherogenic agents. Consistent with this, we found Por-HDL-NPs reduced plaque area *in vivo* in early-stage and unstable atherosclerosis models. In addition to Por-HDL-NPs *in vitro* anti-inflammatory and cholesterol efflux effects, we observed reductions in circulating and aortic monocytes, and aortic arch *Rela* expression. These atheroprotective effects occurred independently plasma cholesterol changes, highlighting their potential to add benefit on top of lipid-lowering.

In conclusion, Por-HDL-NPs exhibit therapeutic and diagnostic properties in atherosclerotic CVD. *In vivo*, Por-HDL-NPs longitudinally detect plaque growth by PET and are detected localized to plaques and plaque macrophages via fluorescence. *In vitro* they are internalized by macrophages and promote cholesterol efflux. Por-HDL-NPs exhibit anti-inflammatory properties and suppress p65-NF-κB, largely independent of cholesterol efflux and SR-BI. Por-HDL-NPs reduce plaque size in early-stage and unstable tandem stenosis models of atherosclerosis, and reduce circulating monocytes and aortic arch *Rela* – potential mechanisms. This is the first characterization of the theranostic properties of Por-HDL-NPs in atherosclerosis. Our findings have significant implications for the use of Por-HDL-mimetic nanoparticles that simultaneously improve diagnosis and prevent plaque development.

## Acknowledgements

The authors acknowledge funding from the Centre of Excellence for Nanoscale BioPhotonics, through the Australian Research Council (ARC, CE140100003) and the National Health and Medical Research Centre (C.A.B and G.Z., Ideas grant: APP1184571). C.A.B. is the recipient of the Lin Huddleston Heart Foundation Fellowship. V.Z. received a University of Adelaide Research training Program Stipend.

---

**Supplemental Figure S1:**
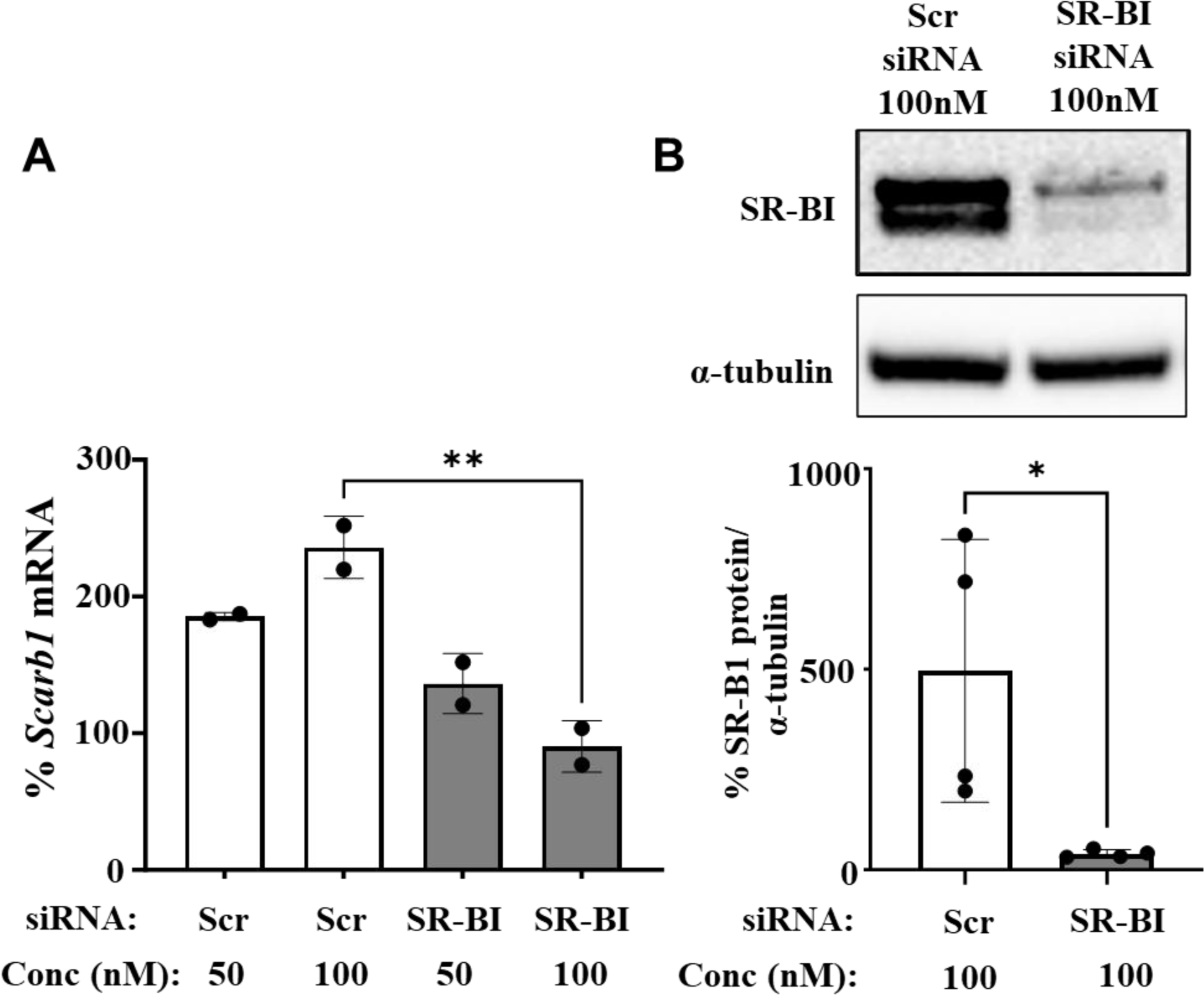
siRNA knockdown of SR-BI in cultured iBMDM macrophages. iBMDMs were transfected with Scrambled (Scr) or SR-BI siRNA (50 and 100 nM). After a 48h incubation iBMDMs were harvested for: (A) RT-qPCR analysis of *Scarb1* (*Sr-b1*) mRNA levels on isolated iBMDM RNA (n=2) or (B) Western blot analyses of SR-BI protein on iBMDM lysates (n=4). Western blot image (top panel) showing SR-BI and α-tubulin as the loading control. Mean ± SD. \**P*<0.05, \*\**P*<0.01.

**Supplemental Figure S2:**
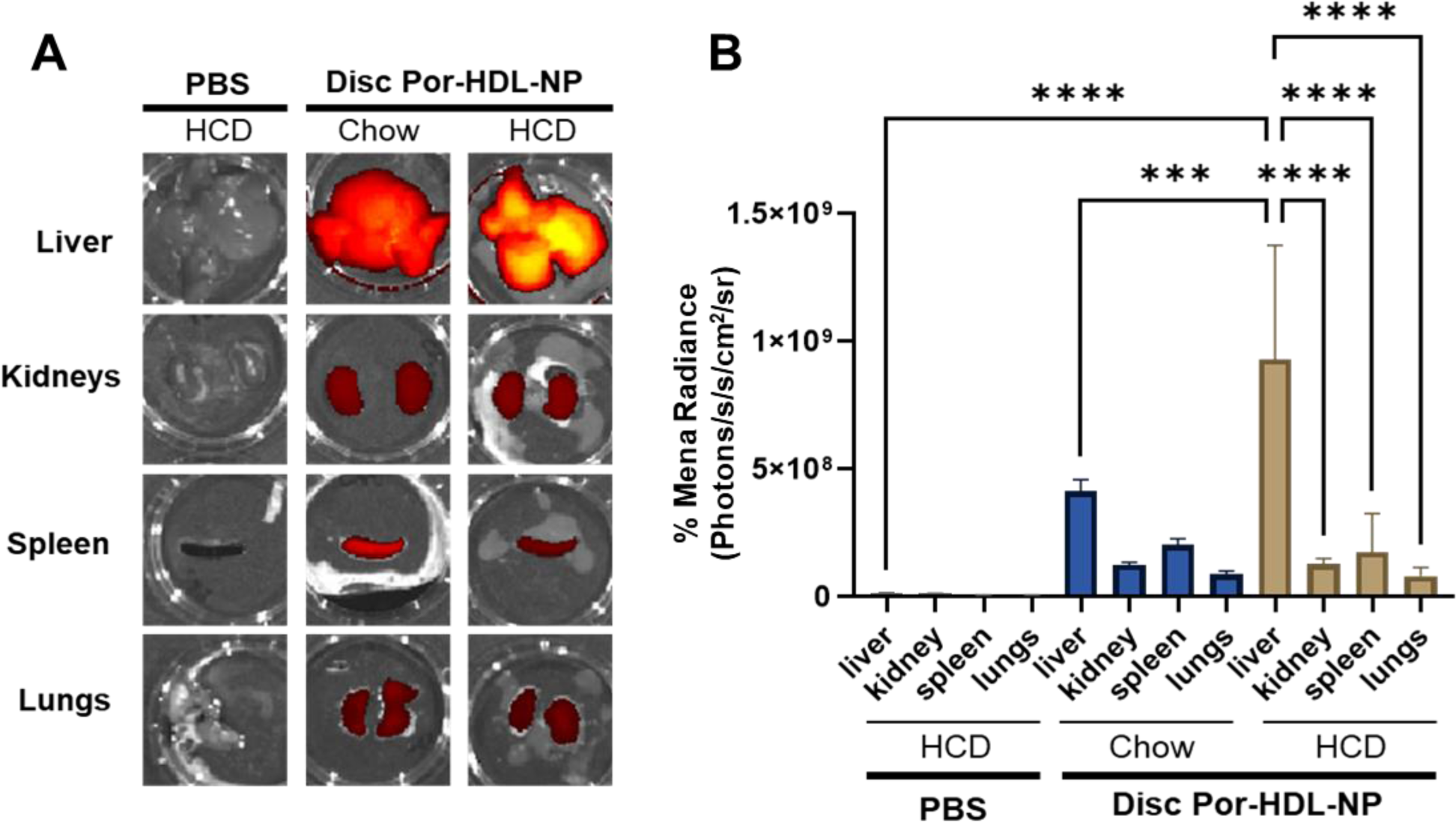
Biodistribution of Por-HDL-NPs in organs of chow and HCD fed *Apoe^-/-^* mice detected *ex vivo* using fluorescence imaging. (A) Representative images of excised liver, kidneys, spleen and lungs from *Apoe^-/-^*mice fed either a chow or HCD collected 24h following intraperitoneal injection of PBS or discoidal 30 mol % Por-HDL-NPs. (B) Fluorescence quantified from IVIS images of excised organs expressed as mean radiance (Photons/s/cm^2^/sr). n=1-4 mice/group (PBS, n=1; Por-HDL-NP Chow, n=1; Por-HDL-NP HCD, n=4). Fluorescence images of Por-HDL-NP chow and HCD organs were captured side-by-side in a single image with the IVIS camera. Data expressed as Mean ± SD. \*\*\**P*<0.001, \*\*\*\**P*<0.0001 by one-way ANOVA with Tukey’s multiple comparisons. IVIS: *In vivo* imaging system; s: second, sr: steradian. Por: Por-HDL-NP.

**Supplemental Figure S3:**
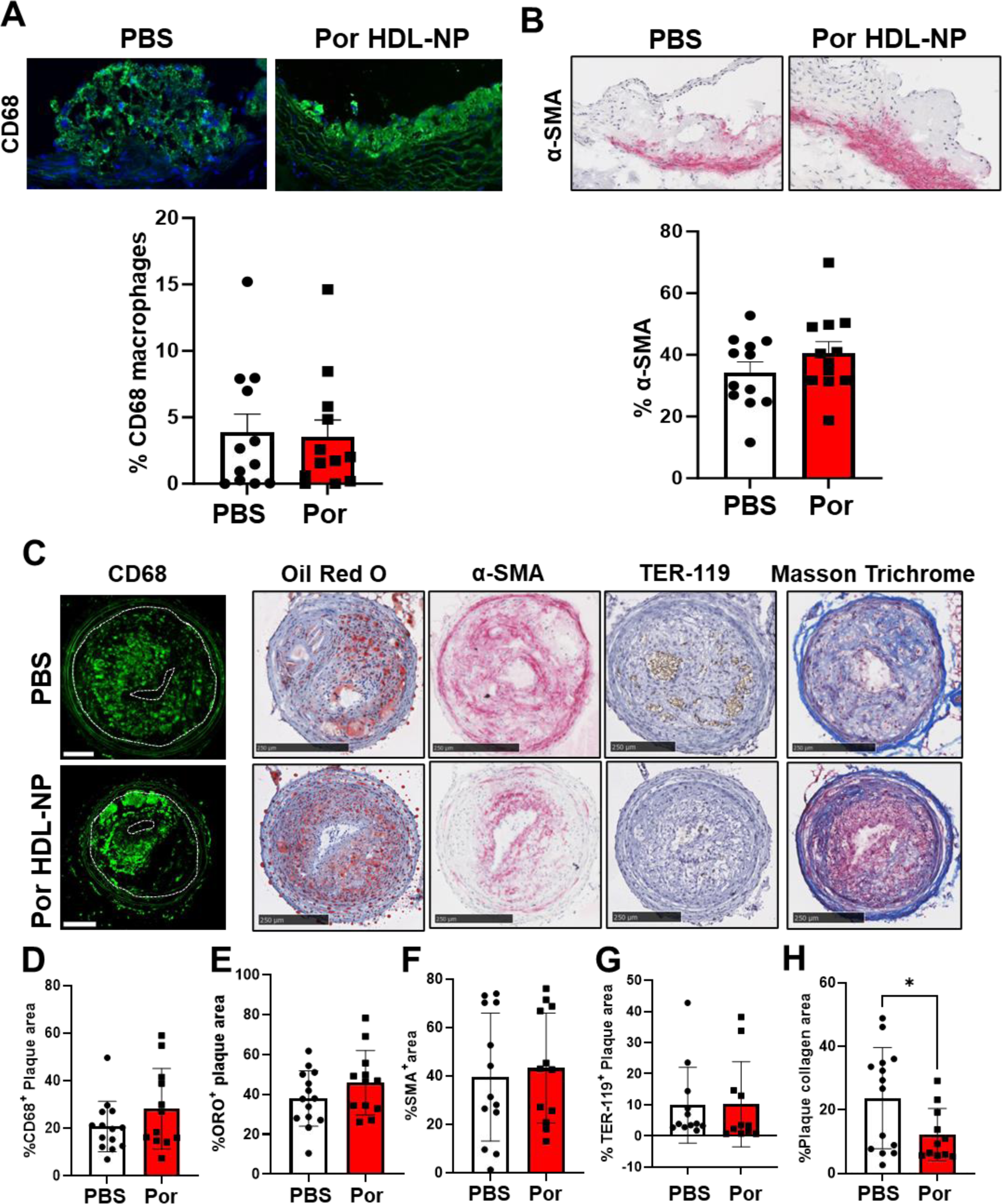
Limited compositional changes observed in plaques from Por-HDL-NP treated mice. Six-week-old *Apoe*^-/-^ mice were fed HCD for 6 weeks to develop early-stage plaque in the aortic sinus and received PBS or Por-HDL-NP infusions three times weekly. Analysis and representative microscopy images of aortic sinus sections assessed for: (A) CD68^+^ macrophages and (B) α-SMA^+^ smooth muscle cells, calculated as a % of total plaque area. (C) After 6 weeks of HCD, *Apoe*^-/-^ mice received tandem stenosis carotid artery ligation surgery. The mice were then infused with either PBS or Por-HDL-NP on alternate days for a further 7 weeks. Representative images of fluorescence and light microscopy of carotid artery sections to detect: (D) CD68^+^ macrophages, (E) Oil Red O lipid, (F) α-SMA^+^ smooth muscle cells, (G) TER-119^+^ erythrocytes and (H) collagen (stained blue in Masson’s Trichrome). Data expressed as mean ± SD (n=12-14/group). \**P*<0.05. Scale bar: CD68 = 200 µm; Oil Red O, TER-119 and Masson’s Trichrome = 500 µm; α-SMA = 250 µm. Por: Por-HDL-NP.

**Supplemental Table S1.**
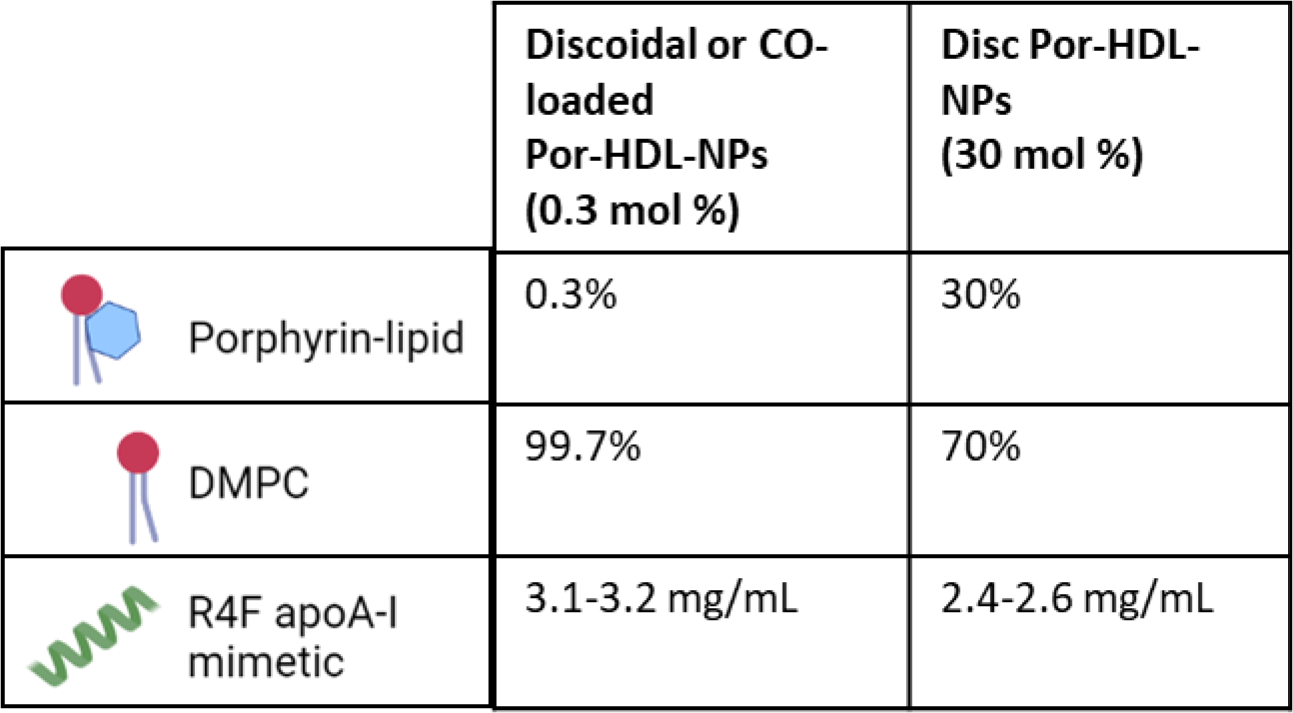
Por-HDL-NPs with compositions of porphyrin-lipid, DMPC phospholipid (% of total moles of lipid per particle) outlined for each nanoparticle formulation. 1,2-Dimyristoyl-sn-glycero-3-phosphocholine, DMPC.

|  | Discoidal or CO-loaded<br>Por-HDL-NPs<br>(0.3 mol %) | Disc Por-HDL-NPs<br>(30 mol %) |
| --- | --- | --- |
| Porphyrin-lipid | 0.3% | 30% |
| DMPC | 99.7% | 70% |
| R4F apoA-I<br>mimetic | 3.1-3.2 mg/mL | 2.4-2.6 mg/mL |

**Supplemental Table S2.**
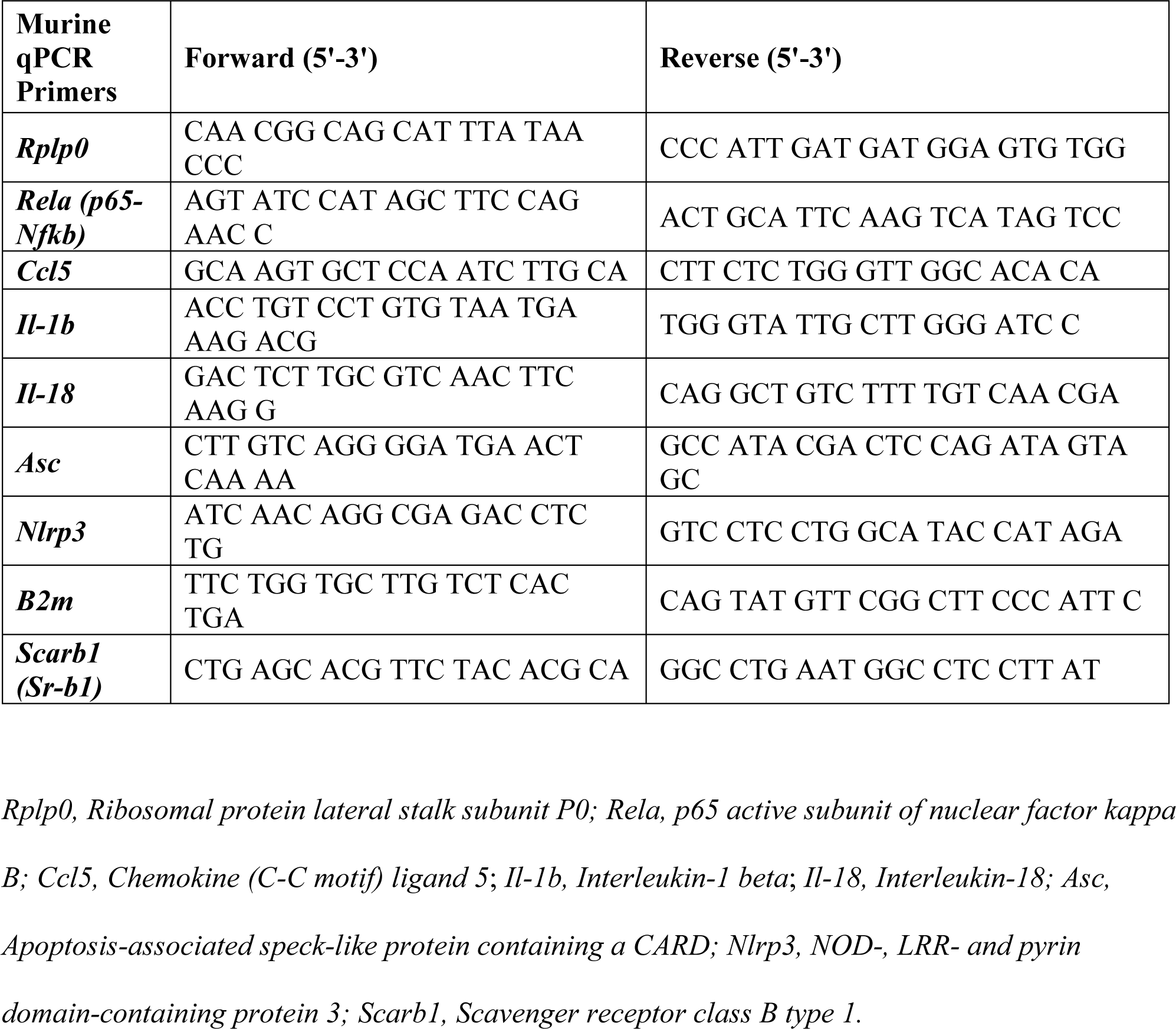
RT-qPCR primer sequences used to measure mRNA levels of inflammatory genes in murine iBMDMs and aortic arches.

**Supplemental Table S3.** Flow cytometry antibodies.

**Flow cytometry antibodies**
| Antibody/ Stain | Catalog #, Manufacturer | 841 |
| --- | --- | --- |
| Anti-Actin, α-Smooth Muscle – FITC, Mouse monoclonal | F3777, Sigma-Aldrich Inc., MO, USA | 842 |
|  |  | 843 |
| APC-Cy<sup>TM</sup>7 Rat Anti-CD11b | 557657, BD Biosciences, NJ, USA | 844 |
| Brilliant Violet 510<sup>TM</sup> anti-mouse Ly-6G/Ly-6C (Gr-1) Antibody | 108438, Biolegend, CA, USA | 845 |
| Brilliant Violet 785<sup>TM</sup> anti-mouse F4/80 | 123141, Biolegend, CA, USA | 846 |
| BV421 Rat Anti-Mouse CD45 | 563890, BD Biosciences, NJ, USA | 847 |
| BV421 Rat Anti-Mouse CD86 | 564198, BD Biosciences, NJ, USA | 848 |
| FITC anti-mouse CD192 (CCR2) Antibody | 150607, Biolegend, CA, USA |  |
| PE Rat Anti-CD11b | 553311, BD Biosciences, NJ, USA | 849 |
| PE/Cyanine7 anti-mouse CD206 (MMR) | 141720, Biolegend, CA, USA | 850 |
| PE-Cy<sup>TM</sup>7 Rat Anti-Mouse Ly-6G | 560601, BD Biosciences, NJ, USA | 851 |
| PerCp-Cy<sup>TM</sup>5.5 Rat Anti-mouse CD31 | 562861, BD Biosciences, NJ, USA | 852 |
| PerCP-Cy<sup>TM</sup>5.5 Rat Anti-Mouse TER-119/Erythroid Cells | 560512, BD Biosciences, NJ, USA | 853 |

**Supplemental Table S4.**

| Plasma Lipids | PBS | Por HDL-NP | P value |
| --- | --- | --- | --- |
| Total Cholesterol (mmol/L) | 5.4 ± 1.2 | 5.3 ± 1.2 | 0.84 |
| LDL Cholesterol (mmol/L) | 3.7 ± 1.5 | 3.1 ± 1.6 | 0.44 |
| HDL Cholesterol (mmol/L) | 3.1 ± 2.4 | 3.1 ± 1.8 | 0.99 |
| Triglycerides (mmol/L) | 5.9 ± 2.2 | 4.9 ± 1.5 | 0.22 |
| Body weight (g) | 26.5 ± 0.4 | 28.1 ± 0.6* | 0.04 |

Plasma lipids and weights of *Apoe*<sup>-/-</sup> mice infused 3 x weekly from 6 weeks of age with PBS or Por-HDL-NPs intraperitoneally for 6 weeks whilst on HCD. Plasma lipids were measured by enzymatic assay. n = 12. Mean ± SD.

**Supplemental Table S5.**
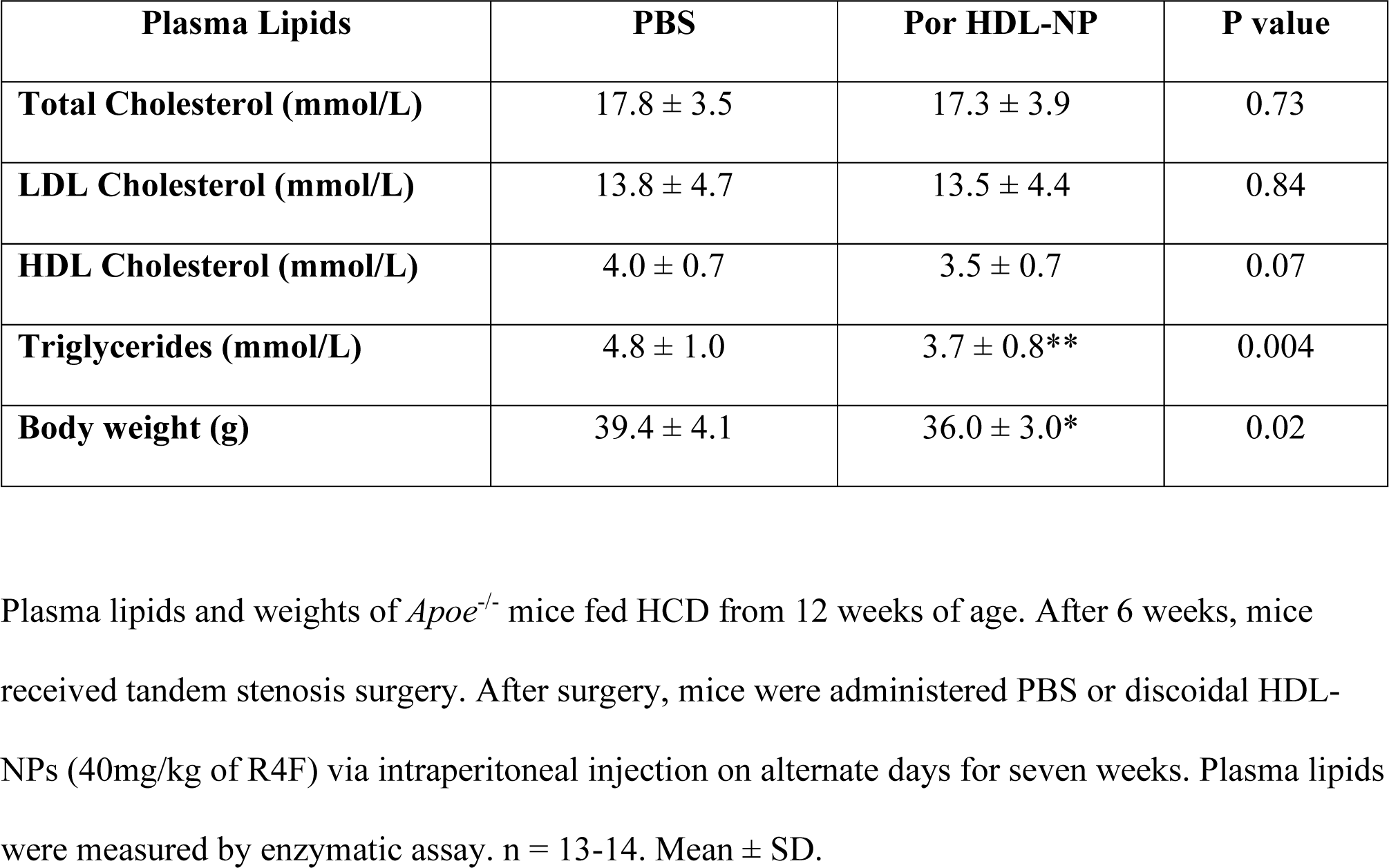

## SUPPLEMENTAL METHODS

### Porphyrin HDL mimetic nanoparticle synthesis

Porphyrin HDL mimetic nanoparticles (Por-HDL-NPs) consist of an outer shell of the phospholipid 1,2-Dimyristoyl-sn-glycero-3-phosphocholine (DMPC), porphyrin-lipid and R4F. R4F is an apoA-I alpha helical mimetic peptide (Ac-<u>F</u>AEK<u>F</u>KEAVKDY<u>F</u>AK<u>F</u>WD). The porphyrin lipid was formed by conjugation of pyropheophorbide-a (porphyrin) and single chain phospholipid 1-palmitoyl-2-hydroxy-sn-glycero-3-phosphocholine (16:0 Lyso PC) as previously described^14,15^. Discoidal and cholesterol oleate (CO)-loaded^37^ Por-HDL-NPs containing 0.3% porphyrin-lipid of the total moles of lipid (0.3 mol %) were used for *in vitro* studies. For imaging studies, the highest porphyrin-lipid content consisted of the discoidal 30 mol % Por-HDL-NPs which contain 30% porphyrin-lipid of the total moles of lipid. Nanoparticle compositions are summarized in Table S1.

### Reconstituted high-density lipoprotein (rHDL)

Apolipoprotein A-I (ApoA-I), the main protein component of native HDL, was isolated from pooled donated plasma samples from healthy humans obtained from the Australian Red Cross (Supply Agreement 14-02NSW-04) by ultracentrifugation and anion-exchange chromatography, as described previously^38–41^. Discoidal rHDL was prepared by complexing purified lipid free apoA-I with phospholipid 1-palmitoyl-2-linoleoyl-phosphatidycholine (PLPC) at an initial PLPC: apoA-I ratio of 100:1. The rHDL was filter sterilised prior to use in the cholesterol efflux experiments. The final apoA-I concentration was determined using the BCA assay.

### Cell culture

Immortalized murine bone marrow derived macrophages (iBMDMs) were kindly provided by Dr Ashley Mansell (Hudson Institute of Medical Research, Victoria, Australia). iBMDMs were maintained in low glucose DMEM (D5523, Sigma) supplemented with 10% FBS (Sigma) in a 37°C, 5% CO_2_ incubator. Cells were seeded 24h prior to treatments.

### Transmission Electron Microscopy

Transmission electron microscopy (TEM) data acquisition and analysis was conducted as per Rajora *et al.*^37^. Briefly, transmission electron microscopy was conducted with a FEI Tecnai T20 electron microscope at the Nanoscale Biomedical Imaging Facility, Peter Gilgan Centre for Research and Learning. Samples were diluted 20-50x in ddH_2_O and loaded onto charged carbon coated grids. The grids were stained with 2% uranyl acetate negative stain and imaged at direct magnification of 60,000 to 100,000x.

### Real time-qPCR

Total RNA was extracted from iBMDMs and murine aortic arches with TRI-reagent (Sigma-Aldrich). RNA concentration was quantified using nanodrop spectrophotometer (Thermofisher Scientific) with the purity of the samples assessed using the absorbance ratio (A260/A280 ∼2.0 for RNA). Then 1400-1845 ng (iBMDMs) or 500ng (aortic arch) of total RNA was reverse transcribed to cDNA using iScript Reverse Transcriptase Supermix (Biorad). Primers for inflammatory and housekeeper genes (Table S2) were used to measure gene expression changes by qPCR. Gene expression was calculated using the ^ΔΔ^*Ct* method referenced to the housekeeper gene *Rplp0* or *Beta-2 microglobulin* (*B2m)*.

### Histology

#### Embedding and sectioning of aortic sinus and tandem stenosis carotid segment I

A transverse cut was made through the entire heart positioned perpendicular to the base of the two atria and ∼1 mm below and proximal side embedded in optimal cutting temperature (OCT) compound (TissueTek). For the tandem stenosis model^20^, Segment I of the carotid (region below proximal suture) was embedded upright in OCT. To section the aortic sinus, embedded hearts were trimmed until the first leaflet emerged and sections collected from this point. At least sixty 7 µm sections of the OCT-embedded aortic sinus were collected continuously on a cryostat (Leica CM3050S) and mounted onto microscope slides (Superfrost Plus). Additionally, the carotid artery segment of the tandem stenosis model that develops unstable plaque, tandem stenosis Segment I, was serially sectioned at 6 µm^20^. Segment I was sectioned from the proximal suture for a length of approximately 2.3 mm at ∼96 µm intervals.

#### Hematoxylin and Eosin (H&E) staining

Fresh frozen OCT-embedded sections were fixed with 10% neutral buffered formalin then stained using a standard H&E staining protocol. Briefly, sections were stained in Mayer’s Hematoxylin, 0.3% acid ethanol, Scott’s tap water and Eosin. Sections were dehydrated and mounted in Dibutylphthalate Polystyrene Xylene (DPX, Sigma-Aldrich Inc., MO, USA).

#### Masson’s Trichrome staining

Fresh frozen OCT-embedded sections were fixed in 4% PFA before incubation in Bouin’s fluid overnight at room temperature. Masson’s trichrome staining was conducted using a Trichrome Stain Kit following the manufacturer’s instructions (ab150686, Abcam). Sections were dehydrated and mounted in DPX.

#### Oil red O staining

Fresh frozen OCT-embedded sections were fixed in 10% neutral buffered formalin. Sections were washed with 60% v/v isopropanol then stained in Oil red O solution (0.6% w/v oil red O, 60% v/v isopropanol) and differentiated in 60% v/v isopropanol. Sections were mounted with Aquatex (Merck-Millipore).

#### Alpha-smooth muscle actin (α-SMA) immunohistochemistry

OCT-embedded sections were fixed in 2% PFA for 10 min. Sections were blocked with 10% goat serum (Sigma-Aldrich Inc., MO, USA), incubated with α-SMA conjugated to alkaline phosphatase antibody (clone 1A4, A5691, Sigma-Aldrich Inc., MO, USA) diluted 1:100, then alkaline phosphatase development with Vector Red substrate kit (Vector Laboratories, Burlingame, CA, USA) following the manufacturer’s instructions. Sections were dehydrated and mounted in DPX.

#### CD68 immunofluorescence staining

Sections were fixed with 2% PFA for 10 min and then washed with 0.1 M glycine solution. Sections were blocked with 5% goat serum and incubated with anti-CD68 (rat monoclonal, Clone FA-11, IgG2a, MCA1957GA, Biorad) diluted 1:250 or Rat IgG2a κ Isotype Control (BD Biosciences) diluted 1:1250 then Donkey anti-Rat IgG (H+L) Alexa Fluor 488 (AF-488) secondary antibody (A-21208, Invitrogen) diluted 1:2000. Sections were mounted with VECTASHIELD® Antifade Mounting Medium with DAPI (Vector Laboratories).

#### TER-119 immunohistochemistry

Fresh frozen OCT-embedded sections were fixed in ice-cold acetone. Sections were incubated in 3% v/v H_2_O_2,_ blocked with 10% normal horse serum then avidin and biotin. Sections were incubated with TER-119 biotinylated rat anti-mouse monoclonal antibody (Thermofisher, MA, USA) diluted 1:400. TER-119 was detected with Vectastain Elite ABC HRP kit according to manufacturer’s instructions (Vector Laboratories, CA, USA). Sections were dehydrated and mounted in DPX.

#### Microscopy

Stained sections were imaged under the Axiolab microscope attached to a camera (Zeiss) or imaged with Nanozoomer digital slide scanner (Hamamatsu). Fluorescence in aortic sinus sections was detected using the 20x objective on an Axio Scan.Z1 slide scanner microscope (Zeiss) with CD68-AF488 staining and porphyrin-lipid detected on the AF488 and Cy5 fluorescence channels respectively. Fluorescence in segment I in the carotid artery of the tandem stenosis model was detected using the 10x objective on the Eclipse NiE fluorescence microscope (Nikon) with CD68-AF488 staining and porphyrin-lipid detected on the FITC and Cy5 fluorescence channels respectively.

#### Histological image analysis

Image analysis of tissue sections was performed using Image Pro-Premier 9.2 (Media Cybernetics) software. For the aortic sinus, three H&E sections spanning the aortic sinus that contained three leaflets were analyzed per animal and expressed as average total lesion area. The area of histological staining from a mid-point section was calculated as a percentage of the selected total lesion area (% of plaque area) for histological, immunochemical and immunofluorescence analyses.

For tandem stenosis Segment I, plaque/lesion area was measured in H&E-stained sections in three separate regions spanning the length of the vessel at 0-384 μm, 768-1152 μm and 1920-2304 μm from the proximal suture. Four sections 96 µm apart in the mid-point region of Segment I (approx. 768-1152 μm from proximal suture) were assessed for histological and immunochemical analyses.

#### Plasma lipid analysis

Triglycerides, total cholesterol, LDL-cholesterol and HDL-cholesterol were measured enzymatically in plasma using commercially available kits (Wako diagnostics). For triglycerides, 2 µL of plasma was using for the assessment, according to manufacturer’s instructions, with color change to blue measured at 600 nm absorbance. The same protocol was followed for total cholesterol, but the plasma was first diluted 1:1 with distilled water. To determine the HDL-cholesterol concentration, 20 µL of plasma was mixed with 20 µL of precipitation reagent from HDL-C kit (Wako diagnostics), then incubated for 10 min at room temperature before centrifugation at 3000 rpm for 15 min. 10 µL of the supernatant was assayed with the total cholesterol kit to determine HDL-C concentration. LDL-C concentration was calculated by subtracting HDL-C concentrations from total cholesterol concentrations.

## References

1. World Health Organisation, Cardiovascular diseases (CVDs). https://www.who.int/en/news-room/fact-sheets/detail/cardiovascular-diseases-(cvds). 2021.

2. Fernández-Friera L, Fuster V, López-Melgar B, Oliva B, García-Ruiz JM, Mendiguren J, Bueno H, Pocock S, Ibáñez B, Fernández-Ortiz A, Sanz J. Normal LDL-Cholesterol Levels Are Associated With Subclinical Atherosclerosis in the Absence of Risk Factors. Journal of the American College of Cardiology. 2017;70:2979–2991. doi: 10.1016/j.jacc.2017.10.024

3. Duivenvoorden R, Tang J, Cormode DP, Mieszawska AJ, Izquierdo-Garcia D, Ozcan C, Otten MJ, Zaidi N, Lobatto ME, van Rijs SM, et al. A statin-loaded reconstituted high-density lipoprotein nanoparticle inhibits atherosclerotic plaque inflammation. Nature communications. 2014;5:3065. doi: 10.1038/ncomms4065

4. Flores AM, Hosseini-Nassab N, Jarr K-U, Ye J, Zhu X, Wirka R, Koh AL, Tsantilas P, Wang Y, Nanda V, et al. Pro-efferocytic nanoparticles are specifically taken up by lesional macrophages and prevent atherosclerosis. Nature Nanotechnology. 2020;15:154–161. doi: 10.1038/s41565-019-0619-3

5. Chen W, Schilperoort M, Cao Y, Shi J, Tabas I, Tao W. Macrophage-targeted nanomedicine for the diagnosis and treatment of atherosclerosis. Nature Reviews Cardiology. 2021. doi: 10.1038/s41569-021-00629-x

6. Zhang Y, Koradia A, Kamato D, Popat A, Little PJ, Ta HT. Treatment of atherosclerotic plaque: perspectives on theranostics. 2019;71:1029–1043. doi: 10.1111/jphp.13092

7. Binderup T, Duivenvoorden R, Fay F, van Leent MMT, Malkus J, Baxter S, Ishino S, Zhao Y, Sanchez-Gaytan B, Teunissen AJP, et al. Imaging-assisted nanoimmunotherapy for atherosclerosis in multiple species. Science Translational Medicine. 2019;11:eaaw7736. doi: 10.1126/scitranslmed.aaw7736

8. Mulder WJM, van Leent MMT, Lameijer M, Fisher EA, Fayad ZA, Pérez-Medina C. High-Density Lipoprotein Nanobiologics for Precision Medicine. Accounts of Chemical Research. 2018;51:127–137. doi: 10.1021/acs.accounts.7b00339

9. Pérez-Medina C, Binderup T, Lobatto ME, Tang J, Calcagno C, Giesen L, Wessel CH, Witjes J, Ishino S, Baxter S, et al. In Vivo PET Imaging of HDL in Multiple Atherosclerosis Models. JACC: Cardiovascular Imaging. 2016;9:950–961. doi: 10.1016/j.jcmg.2016.01.020

10. Chen J, Zhang X, Millican R, Creutzmann JE, Martin S, Jun H-W. High density lipoprotein mimicking nanoparticles for atherosclerosis. Nano Convergence. 2020;7:6. doi: 10.1186/s40580-019-0214-1

11. Rye K-A, Barter PJ. Cardioprotective functions of HDLs. Journal of Lipid Research. 2014;55:168–179. doi: 10.1194/jlr.R039297

12. Rajora MA, Lou JWH, Zheng G. Advancing porphyrin’s biomedical utility via supramolecular chemistry. Chemical Society Reviews. 2017;46:6433–6469. doi: 10.1039/C7CS00525C

13. Liu TW, MacDonald TD, Shi J, Wilson BC, Zheng G. Intrinsically Copper-64-Labeled Organic Nanoparticles as Radiotracers. Angewandte Chemie International Edition. 2012;51:13128–13131. doi: 10.1002/anie.201206939

14. Lovell JF, Jin CS, Huynh E, Jin H, Kim C, Rubinstein JL, Chan WCW, Cao W, Wang LV, Zheng G. Porphysome nanovesicles generated by porphyrin bilayers for use as multimodal biophotonic contrast agents. Nature Materials. 2011;10:324. doi: 10.1038/nmat2986

15. Cui L, Lin Q, Jin CS, Jiang W, Huang H, Ding L, Muhanna N, Irish JC, Wang F, Chen J, Zheng G. A PEGylation-Free Biomimetic Porphyrin Nanoplatform for Personalized Cancer Theranostics. ACS nano. 2015;9:4484–4495. doi: 10.1021/acsnano.5b01077

16. Lin Q, Chen J, Ng KK, Cao W, Zhang Z, Zheng G. Imaging the cytosolic drug delivery mechanism of HDL-like nanoparticles. Pharmaceutical research. 2014;31:1438–1449. doi: 10.1007/s11095-013-1046-z

17. Shen W-J, Azhar S, Kraemer FB. SR-B1: A Unique Multifunctional Receptor for Cholesterol Influx and Efflux. Annu Rev Physiol. 2018;80:95–116. doi: 10.1146/annurev-physiol-021317-121550

18. Linton MF, Tao H, Linton EF, Yancey PG. SR-BI: A Multifunctional Receptor in Cholesterol Homeostasis and Atherosclerosis. Trends Endocrinol Metab. 2017;28:461–472. doi: 10.1016/j.tem.2017.02.001

19. Muhanna N, Cui L, Chan H, Burgess L, Jin CS, MacDonald TD, Huynh E, Wang F, Chen J, Irish JC, Zheng G. Multimodal Image-Guided Surgical and Photodynamic Interventions in Head and Neck Cancer: From Primary Tumor to Metastatic Drainage. Clinical cancer research: an official journal of the American Association for Cancer Research. 2016;22:961–970. doi: 10.1158/1078-0432.ccr-15-1235

20. Chen YC, Bui AV, Diesch J, Manasseh R, Hausding C, Rivera J, Haviv I, Agrotis A, Htun NM, Jowett J, et al. A novel mouse model of atherosclerotic plaque instability for drug testing and mechanistic/therapeutic discoveries using gene and microRNA expression profiling. Circ Res. 2013;113:252–265. doi: 10.1161/CIRCRESAHA.113.301562

21. Rye K-A, Clay MA, Barter PJ. Remodelling of high density lipoproteins by plasma factors. Atherosclerosis. 1999;145:227–238. doi: 10.1016/S0021-9150(99)00150-1

22. Gisterå A, Ketelhuth DFJ, Malin SG, Hansson GK. Animal Models of Atherosclerosis– Supportive Notes and Tricks of the Trade. Circulation Research. 2022;130:1869–1887. doi: 10.1161/CIRCRESAHA.122.320263

23. Sanchez-Gaytan BL, Fay F, Lobatto ME, Tang J, Ouimet M, Kim Y, van der Staay SE, van Rijs SM, Priem B, Zhang L, et al. HDL-mimetic PLGA nanoparticle to target atherosclerosis plaque macrophages. Bioconjug Chem. 2015;26:443–451. doi: 10.1021/bc500517k

24. Sei YJ, Ahn J, Kim T, Shin E, Santiago-Lopez AJ, Jang SS, Jeon NL, Jang YC, Kim Y. Detecting the functional complexities between high-density lipoprotein mimetics. Biomaterials. 2018;170:58–69. doi: 10.1016/j.biomaterials.2018.04.011

25. Marrache S, Dhar S. Biodegradable synthetic high-density lipoprotein nanoparticles for atherosclerosis. Proceedings of the National Academy of Sciences. 2013;110:9445. doi: 10.1073/pnas.1301929110

26. Huynh E, Zheng G. Organic Biophotonic Nanoparticles: Porphysomes and Beyond. IEEE Journal of Selected Topics in Quantum Electronics. 2014;20:27–34. doi: 10.1109/JSTQE.2013.2289971

27. VanderLaan PA, Reardon CA, Getz GS. Site Specificity of Atherosclerosis. *Arteriosclerosis*, Thrombosis, and Vascular Biology. 2004;24:12–22. doi: 10.1161/01.ATV.0000105054.43931.f0

28. Ley K, Miller YI, Hedrick CC. Monocyte and Macrophage Dynamics During Atherogenesis. *Arteriosclerosis*, Thrombosis, and Vascular Biology. 2011;31:1506–1516. doi: 10.1161/ATVBAHA.110.221127

29. Song GJ, Kim S-M, Park K-H, Kim J, Choi I, Cho K-H. SR-BI mediates high density lipoprotein (HDL)-induced anti-inflammatory effect in macrophages. Biochemical and Biophysical Research Communications. 2015;457:112–118. doi: 10.1016/j.bbrc.2014.12.028

30. Grebe A, Hoss F, Latz E. NLRP3 Inflammasome and the IL-1 Pathway in Atherosclerosis. Circulation Research. 2018;122:1722–1740. doi: 10.1161/CIRCRESAHA.118.311362

31. Ridker PM, Everett BM, Thuren T, MacFadyen JG, Chang WH, Ballantyne C, Fonseca F, Nicolau J, Koenig W, Anker SD, et al. Antiinflammatory Therapy with Canakinumab for Atherosclerotic Disease. New England Journal of Medicine. 2017;377:1119–1131. doi: 10.1056/NEJMoa1707914

32. Libby P. Interleukin-1 Beta as a Target for Atherosclerosis Therapy: Biological Basis of CANTOS and Beyond. Journal of the American College of Cardiology. 2017;70:2278–2289. doi: 10.1016/j.jacc.2017.09.028

33. Baldrighi M, Mallat Z, Li X. NLRP3 inflammasome pathways in atherosclerosis. Atherosclerosis. 2017;267:127–138. doi: 10.1016/j.atherosclerosis.2017.10.027

34. Swanson KV, Deng M, Ting JPY. The NLRP3 inflammasome: molecular activation and regulation to therapeutics. Nature Reviews Immunology. 2019;19:477–489. doi: 10.1038/s41577-019-0165-0

35. Gordon T, Castelli WP, Hjortland MC, Kannel WB, Dawber TR. High density lipoprotein as a protective factor against coronary heart disease. The Framingham Study. The American journal of medicine. 1977;62:707–714.

36. Rohatgi A, Westerterp M, von Eckardstein A, Remaley A, Rye K-A. HDL in the 21st Century: A Multifunctional Roadmap for Future HDL Research. Circulation. 2021;143:2293–2309. doi: 10.1161/CIRCULATIONAHA.120.044221

37. Rajora MA, Ding L, Valic M, Jiang W, Overchuk M, Chen J, Zheng G. Tailored theranostic apolipoprotein E3 porphyrin-lipid nanoparticles target glioblastoma Chemical Science. 2017;8:5371-5384. doi: 10.1039/c7sc00732a

38. Tan JTM, Prosser HCG, Dunn LL, Vanags LZ, Ridiandries A, Tsatralis T, Leece L, Clayton ZE, Yuen SCG, Robertson S, et al. High-Density Lipoproteins Rescue Diabetes-Impaired Angiogenesis via Scavenger Receptor Class B Type I. Diabetes. 2016;65:3091–3103. doi: 10.2337/db15-1668

39. Bursill CA, Castro ML, Beattie DT, Nakhla S, van der Vorst E, Heather AK, Barter PJ, Rye KA. High-density lipoproteins suppress chemokines and chemokine receptors in vitro and in vivo. Arterioscler Thromb Vasc Biol. 2010;30:1773–1778. doi: 10.1161/ATVBAHA.110.211342

40. van der Vorst EP, Vanags LZ, Dunn LL, Prosser HC, Rye KA, Bursill CA. High-density lipoproteins suppress chemokine expression and proliferation in human vascular smooth muscle cells. FASEB J. 2013;27:1413–1425. doi: 10.1096/fj.12-212753

41. Tsatralis T, Ridiandries A, Robertson S, Vanags LZ, Lam YT, Tan JTM, Ng MKC, Bursill CA. Reconstituted high-density lipoproteins promote wound repair and blood flow recovery in response to ischemia in aged mice. Lipids in Health and Disease. 2016;15:150. doi: 10.1186/s12944-016-0322-4

